# Resetting of the 24-nt siRNA landscape in rice zygotes

**DOI:** 10.1101/2020.08.31.275958

**Authors:** Chenxin Li, Jonathan I. Gent, Hengping Xu, Hong Fu, Scott D. Russell, Venkatesan Sundaresan

## Abstract

**Background:** The zygote, a totipotent stem cell, constitutes a critical stage of the life cycle of sexually reproducing organisms. It is produced by the fusion of two differentiated cells — the egg and sperm, which in plants have radically different siRNA transcriptomes from each other and from multicellular embryos. Due to technical challenges, the epigenetic changes that accompany the zygotic transition are poorly understood.

**Results:** Here, we characterized the small RNA transcriptome of rice zygotes. We found widespread redistribution of 24-nt siRNAs relative to gametes, including absence of sperm signature siRNAs, reduction at egg signature siRNA loci, and upregulation at seedling signature siRNA loci. Loci with reduced siRNAs in zygote relative to egg were gene-distal and heterochromatic, while loci with increased siRNAs relative to egg had a similar genomic distribution to canonical siRNA loci. Although both egg and zygote siRNA loci had higher mCHH level in wildtype than in *drm2* embryo, zygote but not egg siRNA loci were associated with hypermethylation in mature embryo. A small fraction of siRNA loci (~1%) called siren loci accounted for 60% of all siRNAs within zygote siRNA loci, that likely arose from maternal carryover as they had similarly abundant siRNAs in egg; these siren loci were not associated with embryo hypermethylation.

**Conclusions:** Taken together, our results indicate re-distribution of siRNAs in rice zygotes towards the canonical vegetative profile, that are consistent with the initiation of resetting of the gametic epigenome before the first embryonic division.

## Background

Gametes and zygotes constitute critical developmental stages in the life cycle of all sexually reproducing organisms. During fertilization, the egg cell fuses with a sperm cell to form the zygote, which is an undifferentiated and totipotent stem cell that initiates embryogenesis. Flowering plants undergo double fertilization, in which a second sperm cell fuses with the central cell and gives rise to the endosperm, a nutritive tissue that nurtures the developing embryo or germinating seedling [reviewed in (Lord and Russell, 2002)]. In animals, early embryogenesis is controlled by maternal gene products pre-deposited in the egg cell. Depending on the organism, the zygotic genome does not become transcriptionally active until a number of cell divisions have occurred (Tadros and Lipshitz, 2009). Recent studies show that material-to-zygote-transition in flowering plants differs markedly from most animals [reviewed in (Armenta-Medina and Gillmor, 2019)]. In rice zygotes, thousands of genes are upregulated in zygotes, many of which are undetected in the egg cell, consistent with similar observations in zygotes of maize and *Arabidopsis* (Chen et al., 2017; Zhao et al., 2019). Furthermore, zygotic transcription was shown to be required for early embryogenesis (Kao and Nodine, 2019; Zhao et al., 2019). These observations suggest that in angiosperms, unlike most animals, zygotes are transcriptionally active, and zygotic genome activation (ZGA) occurs in the zygote. However, similar to animals, ZGA in plants is gradual. The initial transcriptome of flowering plant zygotes is thus dominated by egg cell RNA carryover, and although newly expressed genes in the zygote are widespread and represent a significant fraction of the zygote transcriptome, their expression levels are relatively low (Anderson et al., 2017; Chen et al., 2017; Zhao et al., 2019).

Along with dynamic changes in gene expression, epigenomic reprogramming has been observed during flowering plant reproduction. In rice and maize, the egg cell is ~10 times larger than sperm in diameter, and thus ~1000 times larger than the sperm cell in volume (Anderson et al., 2013; Kranz et al., 1991), and its chromatin is diffused (Scholten et al., 2002). In contrast, the sperm cell chromatin undergoes global condensation, paralleling animal sperm chromatin in which protamines replace histones (Kimmins and Sassone-Corsi, 2005). A male-germline specific histone H3 variant MGH3 (also termed H3.10) is present in the sperm cell (Borg and Berger, 2015; Okada et al., 2005), following the removal of H3.1 (Borg et al., 2020). H3.10 is resistant to trimethylation at H3K27 (H3K27me3), thus priming the activation of key genes for sperm differentiation and embryogenesis (Borg et al., 2020). Upon karyogamy, H3.10 is removed from the paternal chromatin via a replication independent process (Ingouff et al., 2007). Other histone H3 variants, such as H3.3, are also removed from egg cell chromatin upon karyogamy, followed by loading of newly-synthesized histones, again via a replication independent mechanism (Ingouff et al., 2010). In addition, other cells of both male and female gametophytes in *Arabidopsis* experience global chromatin changes as well. Heterochromatin is decondensed in the central cell, the cell which gives rise to endosperm (Pillot et al., 2010). A similar phenomenon occurs in the pollen vegetative cell, the cell which encapsulates the sperm cells and enables their migration through the style to the ovule (Schoft et al., 2009; Mérai et al., 2014; Hsieh et al., 2016). Relaxation of heterochromatin in the pollen vegetative cell has been reported to produce short interfering RNA (siRNA) that traffic into the sperm cells, and reinforce transposon silencing in the gametes (Slotkin et al., 2009; Calarco et al., 2011; Martínez et al., 2016; Park et al., 2016; Kim et al., 2019). Similarly, it has been proposed that siRNAs traffic from the central cell to the egg cell, as well as from the endosperm into the developing embryo (Hsieh et al., 2009; Ibarra et al., 2012; Martínez and Köhler 2017).

Concomitant with chromatin reprogramming, there is also evidence for changes in DNA methylation during plant reproduction, especially in the context of RNA-directed DNA methylation (RdDM) [reviewed in (Gehring, 2019)]. In plants, RdDM can function in both *de novo* and maintenance DNA methylation [reviewed in Cuerda-Gil, and Slotkin (2016)]. Briefly, 24-nt siRNAs are loaded onto an argonaute protein (AGO), which recruits the DNA methyltransferase Domains Rearranged Methyltransferase2 (DRM2). DRM2 leads to methylation in all sequence contexts, but methylation in the CHH context (mCHH), where H is A, C or T, is a strong indicator of RdDM in both rice and maize (Tan et al., 2016, 2018; Gent et al., 2013), but not in all plants (Zemach et al., 2013). Multiple studies reported that disruption of RdDM leads to a variety of reproductive phenotypes, including aborted embryos (Autran et al., 2011; Grover et al., 2018), arrested pollen (Wang et al., 2020), defective triploid block when the seeds were produced from a 2n maternal × 4n paternal cross (Borges et al., 2018; Erdmann et al., 2017; Martínez et al., 2018; Satyaki and Gehring, 2019) and defective floral development (Dorweiler et al., 2000; Moritoh et al., 2012). These observations suggest siRNAs and RdDM are important for normal plant reproduction.

In mammals, it has long been proposed that fusion of two epigenetically distinct gametes presents a challenge in reproduction, and resetting of the epigenome is required for the pluripotent state of the early embryo [reviewed in (Messerschmidt et al., 2014)]. Epigenome reprogramming in mammals includes large-scale erasure of somatic chromatin signatures in germ cell precursors, establishment of sex-specific signatures in the germline, and post-fertilization resetting towards pluripotency [reviewed in (Messerschmidt et al., 2014; Saitou et al., 2012; Tang et al., 2016)]. The functional consequences of epigenomic changes in gametic fate acquisition and subsequent zygotic totipotency in plants are unclear. It is clear, however, that in plants, the majority of DNA methylation is stably transmitted both maternally and paternally [reviewed in (Gehring, 2019)]. In *C. elegans*, siRNAs can serve as carriers of transgenerational epigenetic information, in which siRNAs can be inherited across a few generations [reviewed in (Houri-Zeevi and Rechavi, 2017)]. While multiple changes in siRNA profiles have been observed during plant reproduction (Calarco et al., 2012; Grover et al., 2020; Ibarra et al., 2012; Li et al., 2020; Papareddy et al., 2020; Schoft et al., 2009; Slotkin et al., 2009), transgenerational inheritance of siRNAs, or the lack thereof, has yet to be rigorously demonstrated in plants.

In vegetative tissues such as seedlings, 24-nt siRNAs coincide with mCHH islands - short regions with high CHH methylation - that are enriched around genes and mark the ends of TEs and euchromatin-heterochromatin boundaries (Gent et al., 2013; Li et al., 2015). Hereafter, we refer to such a 24-nt siRNA profile as the canonical siRNA profile, since 24-nt siRNAs are the most abundant length class in most plants [reviewed in Cuerda-Gil, and Slotkin (2016)], including rice gametes (Li et al., 2020, **Fig 1B**). We have previously shown that the siRNA transcriptome is reprogrammed in rice gametes (Li et al., 2020) where siRNA transcriptomes of egg and sperm were distinct from each other in genome-wide distribution, as well as distinct from that of the seedling (**Fig. 1**). The relative magnitude of the egg-borne and sperm-borne contribution of siRNAs to the zygote is unknown. A recent study in *Arabidopsis* revealed that siRNAs from heterochromatic TEs are transiently upregulated during embryogenesis, while siRNAs from euchromatic TEs peak at mature green embryos (Papareddy et al., 2020). However, due to technical challenges due to the limiting amounts of material, currently there are no data available for siRNA transcriptomes before the preglobular embryo stage, and consequently very little is known about the siRNA landscape in plant zygotes. Since siRNA production is influenced by histone modifications and DNA methylation, and siRNAs in turn can direct histone modifications and DNA methylation (Law and Jacobsen, 2010; Matzke and Mosher, 2014; Parent et al., 2021), the siRNA transcriptome is an output and indicator of the epigenome. Given the likely importance of epigenetic reprogramming during plant reproduction and the lack of detailed studies on zygotes, we decided to characterize the small RNA transcriptome of rice zygotes to investigate changes in the small RNA transcriptome that occur soon after fertilization. The differences in genome-wide distribution of siRNAs between gametes and zygote revealed changes that indicate that the siRNA transcriptome initiates a resetting towards the canonical profile before the first cell division, concurrently with zygotic genome activation.

**Fig 1:**
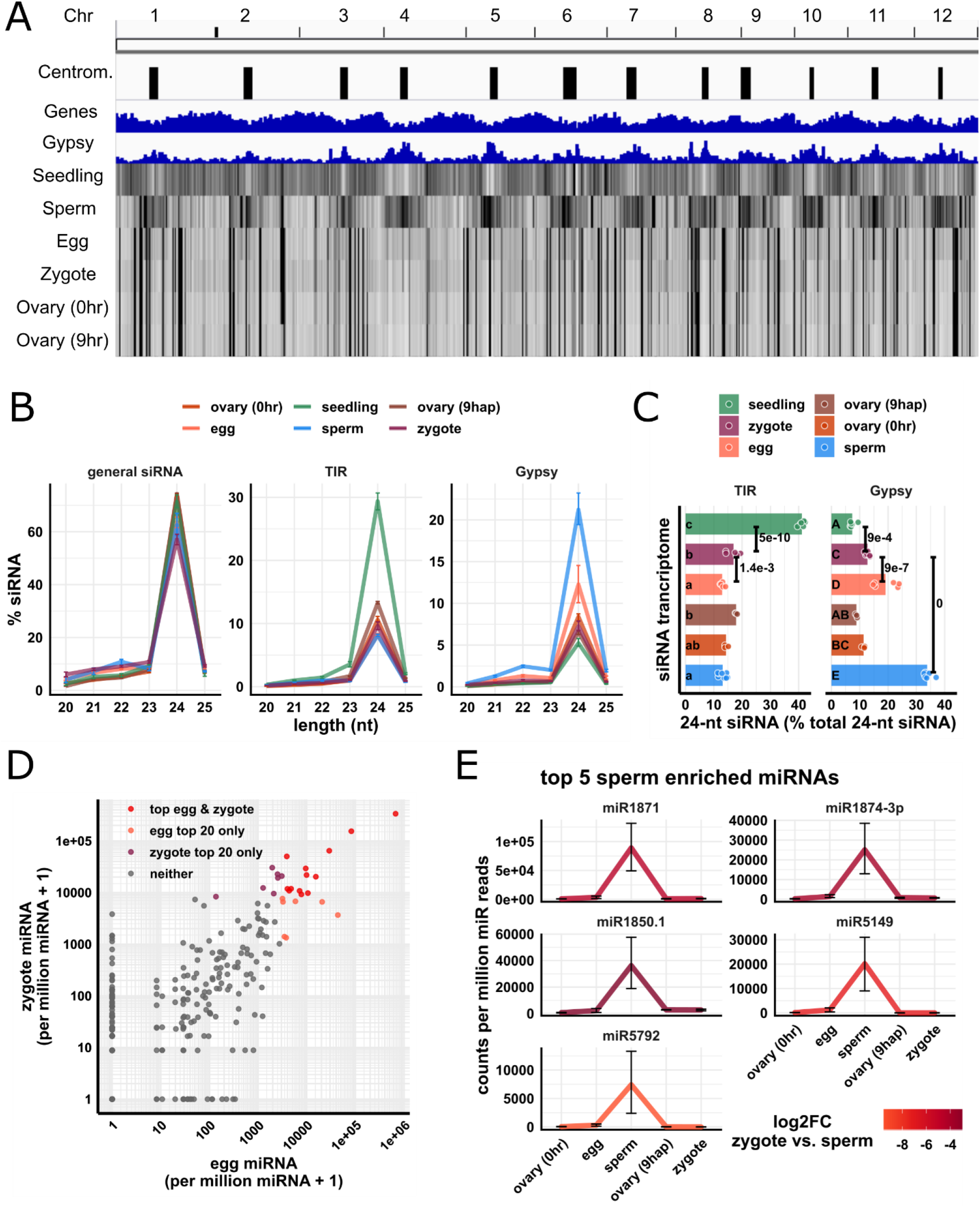
Overall pattern of zygote 24-nt siRNAs is similar to but not identical to egg cell. (A) Heat map showing abundance of 24-nt siRNA across genome at 50-kb resolution. The first three tracks are centromeres [as defined by (Mizuno et al., 2018)], genes, and *Gypsy* retrotransposons. (B) Length profiles of siRNAs. y-axis values are relative to total siRNA reads (20 – 25-nt siRNAs). TIR: terminal inverted repeat transposons, CACTA superfamily excluded. *Gypsy*: *Gypsy* retrotransposons. Error bars are 95% confidence intervals for each cell type. miRNA and phasiRNA are not included in this analysis (**Fig S1A**). (C) Quantification of TIR and *Gypsy* panels in (B). Each data point is an siRNA transcriptome. Bar heights are averages. x-axis values are relative to total 24-nt siRNAs. Letter grouping (α *=* 0.05) and P values are based on Tukey tests. (D) Scatter plot showing miRNA relative abundances in egg and zygote. Each data point is a miRNA. Axes are relative to per million miRNA reads and log10 transformed. ‘top egg & zygote’ refers to intersection of the 20 highest abundant miRNAs in both egg and zygote. (E) Top five sperm enriched miRNAs. Sperm enriched is classified as > 1000 reads per million miRNA reads in sperm and < 500 reads per million miRNA reads in egg. y-axis values are relative to per million miRNA reads. Color code reflects log2FC values for zygote vs. sperm, and negative values indicate higher in sperm. Error bars are 95% confidence intervals for each cell type. See **Fig S1D** for additional examples. Zygote and 9 hap ovary data are from this study, all other data from Li et al., (2020).

## Results

We collected rice zygotes ~9 hours after pollination (hap), which corresponds to the completion of S-phase, just prior to the first zygotic division (Anderson et al., 2017; Ding et al., 2009). We generated small RNA transcriptomes from 6 replicates, with ~50 zygotes in each replicate. As a maternal sporophytic control, we also collected post-fertilization ovary of the same developmental stage as zygote (9 hap) and prepared small RNA transcriptomes from 3 replicates, with 10 ovaries in each replicate. For our analyses, we also included small RNA transcriptome data from rice gametes, pre-anthesis ovary (0 hr ovary) and seedlings (Li et al., 2020). Except where indicated otherwise, siRNAs used for analyses were small RNA reads (20-nt – 25-nt) not overlapping 90% or more of their lengths with known miRNAs [miRBase v22, (Kozomara et al., 2019)], 5S rRNA, tRNA, NOR, or phasiRNA loci [as detected in Li et al., 2020], and multi-mapped reads were included in all analyses unless indicated otherwise (**Fig S1A**).

### The global siRNA pattern in zygote is determined by siRNA transcript carryover from the egg cell, with no detectable signature of sperm cell small RNAs

As we previously reported, the sperm cell has an siRNA pattern complementary to the canonical pattern of vegetative tissues, in which its 24-nt siRNAs are spread out across wide heterochromatic regions, including centromeric tandem repeats. The egg cell and ovary have a pattern different from both sperm and vegetative tissues, in which 24-nt siRNAs are concentrated at discrete loci (**Fig 1A**). We found that in a whole-genome view, the zygote had a similar pattern to the egg cell (**Fig 1A**, zygote vs. egg track). To confirm that the similarity between zygote and egg cell was not due to large numbers of residual unfertilized egg cells in the zygote samples, we performed a control pollination experiment under similar conditions, and we determined that 98 out of 101 pollinated rice florets produced mature seeds, implying that 3% or less of the rice florets were unfertilized (**Supplemental Table 2**, see **Methods** for additional details). Thus, in our zygote samples, unfertilized egg cells might represent at most 3% of the total. We also performed differential expression analyses for miRNAs and detected 14 miRNAs that were lowly expressed in all six replicates of zygote but highly expressed in ovaries of the corresponding developmental stage, i.e., 9 hap (**Fig S1B**). Thus, the similarity between zygote and ovary (**Fig 1A**) is unlikely to be due to small RNA contamination from ovary. A similar analysis was previously used to show that the egg cell samples were also free of pre-fertilization ovary contamination (Li et al., 2020).

We next looked at the length profile of siRNAs in zygotes and compared that with published data from other cell and tissue types (Li et al., 2020). We found that in zygotes, as in all other tissues, 24-nt siRNAs predominated (**Fig 1B**). Since the abundance of siRNAs of other length classes were all relatively low, we focused on 24-nt siRNAs for further analysis. Based on relative abundance patterns, the zygote siRNAs appeared to resemble egg cell siRNAs. Like the egg cell and unlike seedling tissues, the zygote had a lower abundance of siRNAs overlapping terminal inverted repeat (TIR) transposons (PIF/Harbinger, Tc1/Mariner, Mutator, or hAT superfamilies) than seedling (**Fig 1B-C**, seedling vs. zygote *P* = 5e-10, Tukey tests). Like the egg cell and unlike the sperm cell, the zygote had a low abundance of siRNAs overlapping *Gypsy* retrotransposons (**Fig 1B-C**, sperm vs. zygote *P* = 0, Tukey tests). However, we noted that while the zygote and egg cell were similar, there were some clear differences. Zygote had significantly more siRNAs overlapping TIR elements, and significantly less siRNA overlapping *Gypsy* retrotransposons than the egg cell (**Fig 1C**, *P* = 1.4e-3 and *P* = 9e-7 respectively, Tukey tests).

The similarity between egg and zygote siRNA profiles can be explained by carryover from the egg cell, since the egg cell is ~1000-fold larger than the sperm cell by volume (Kranz, Bautor, and Lörz 1991; Anderson et al., 2013; Li et al., 2019). Although 24-nt siRNAs function in the nucleus, 24-nt siRNAs were found primarily in the cytoplasm of whole-plant homogenates (Ye et al., 2012). Thus, we predict that small RNAs already present in the egg cell before fertilization would contribute to much of the siRNAs present in the zygote. This is consistent with previous observations that the 50 most highly expressed genes in egg cell remained as most highly expressed in zygote, whereas the 50 most highly expressed genes in the sperm cell were low expressed in the zygote (Anderson et al., 2017, 2013). Indeed, 13 out of the 20 most abundant miRNAs in egg cells remained among the 20 most abundant miRNAs in zygote (*P* = 3e-14, Exact test, **Fig 1D**). However, the miRNA accumulation patterns were not identical between zygote and egg. 32 miRNAs were detected in the zygote but not in the egg cell (> 50 reads per million miRNA reads in zygote and undetected in egg cell), and 7 miRNAs were detected in the egg cell but not in the zygote (> 50 reads per million miRNA reads in egg cell and undetected in zygote). The presence of 32 miRNAs detected in zygote but not egg cell suggests that ZGA is initiated at miRNA loci at this stage, which would be consistent with the known ZGA of other RNA polymerase II transcripts. Meanwhile, top sperm-enriched miRNAs were very much downregulated in the zygote, consistent with dilution after fertilization (**Fig 1E** and **Fig S1C**). Note that the expression values in the zygote were not used to define these sperm-enriched miRNAs, as we classified sperm-enriched miRNAs relative to egg alone. Specifically, we required >1000 reads per million miRNA reads in sperm, and < 500 reads per million miRNA reads in the egg cell for this classification. The expression values of the full set of expressed miRNA genes [miRBase v22, (Kozomara et al., 2019)] are provided as a complementary transcriptomics resource (**Supplemental Dataset 1**). Taken together, these results imply that sperm small RNAs were diluted by the egg cell cytoplasm, and that much of the siRNAs detected in the zygote were due to carryover from the egg cell.

### Unusual features of siRNA loci with abundant siRNAs in egg, ovary, zygote, and endosperm

Zygote, like egg cell and ovary, had a very uneven siRNA distribution across the genome, where siRNAs appeared to be concentrated at discrete sites, without a clear relationship to gene density (**Fig 1A**). It has been previously reported that rice developing endosperm (7-8 days after fertilization) has a unique siRNA profile in which a small number of loci accounted for the majority of siRNAs (Rodrigues et al., 2013). These siRNA loci were termed siren loci (<u>siR</u>NA in the <u>en</u>dosperm). A similar phenomenon was recently reported in *Brassica rapa* and *Arabidopsis* ovules and endosperm (Grover et al., 2020). The term ‘siren loci’ was also used by Grover et al to describe these loci. To further investigate this phenomenon in zygote as well as egg, ovary, and endosperm, we ranked siRNA loci according to siRNA abundance in each cell type (**Fig 2A**). In endosperm and ovaries (pre- and post-fertilization), ~0.1% (*n* = 73, 213 and 102, respectively) of the siRNA loci accounted for 60% of the total siRNA accumulation in all siRNA loci for each tissue type (**Fig 2A**). Similarly, in egg cell and zygote, ~1% (*n* = 1881 and 1429, respectively) of the siRNA loci accounted for 60% of the total siRNA accumulation in all siRNA loci for each cell type (**Fig 2A**). We call these highly expressing loci siren loci, independently of siRNAs in endosperm. In fact, the siren loci in rice ovaries, egg, and zygote showed little correlation with the siren loci reported in rice endosperm, at least for the specific endosperm stage described, i.e., 7-8 days after fertilization (Li et al., 2020; Rodrigues et al., 2013). Importantly, egg siren loci were stably expressed between egg and zygote, without a significant decrease after fertilization (**Fig 2B**), which likely explains the overall similarity between egg and zygote (**Fig 1A** and **Fig 3E**, see below).

**Fig 2:**
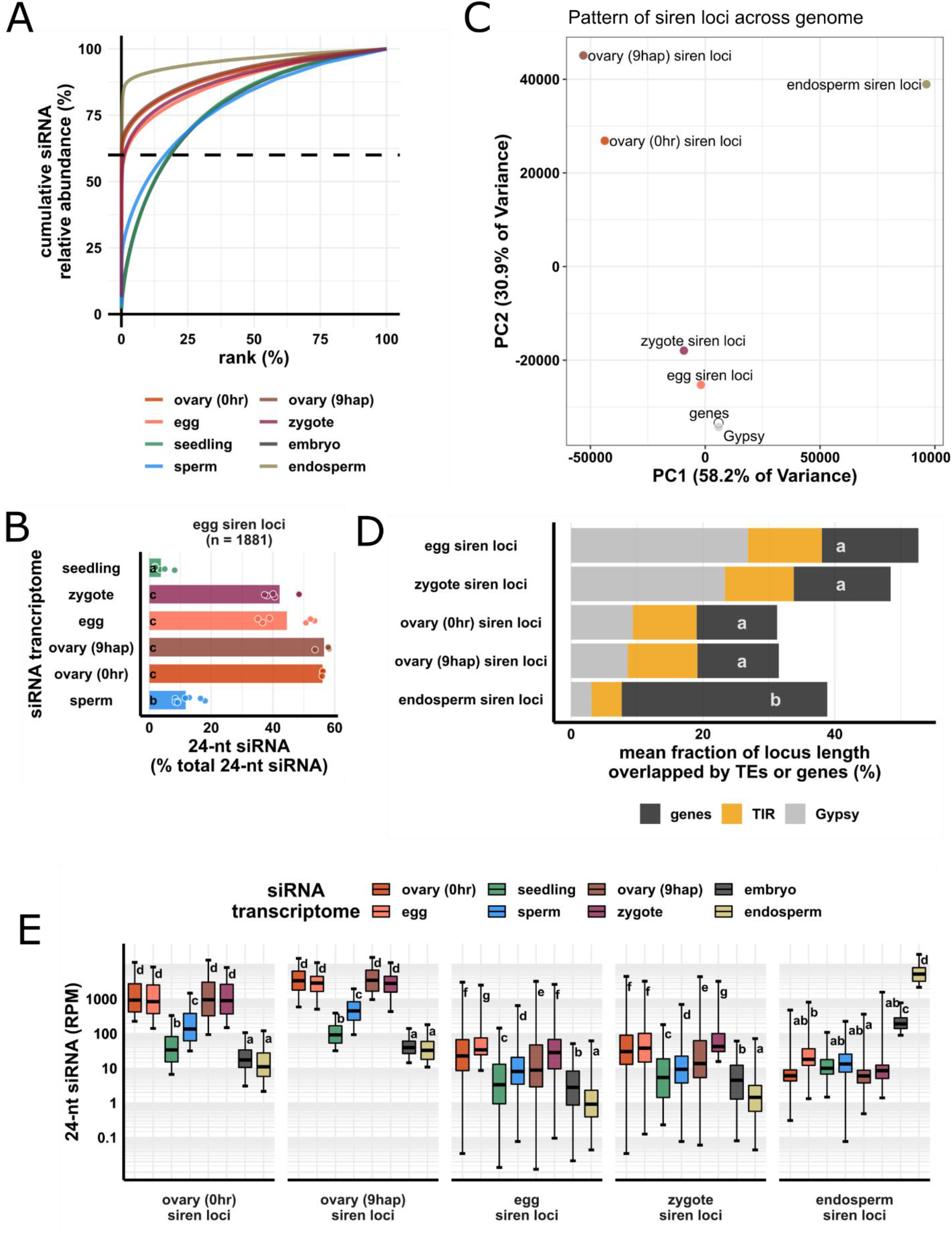
Zygote siren loci are similar to siren loci detected in ovary and egg cell and stably expressed between egg and zygote but dissimilar to siren loci detected in endosperm. (A) x-axis is the rank order of siRNA loci. siRNA loci with highest siRNA abundances are ranked first. y-axis is cumulative relative abundance of siRNA in all siRNA loci. Axis values are scaled between 0 and 100%. 0.1% of siRNA loci accounted for 60% of siRNA reads in all siRNA loci in endosperm and ovary. 1% of siRNA loci accounted for 60% of siRNA reads in all siRNA loci in egg and zygote. (B) Bar plot showing relative abundances of 24-nt siRNA at egg siren loci. Each data point is an siRNA transcriptome. Bar heights are averages. x-axis values are relative to total 24-nt siRNAs. (C) Principal component plot for siren loci distribution across the genome. Distributions are evaluated at 50-kb resolution across the genome. Each data point is the distribution of a siren loci category. (D) Stacked bar plots showing mean fraction of locus length overlapped by TEs or genes. TIR: terminal inverted repeat transposons, CACTA superfamily excluded. *Gypsy*: *Gypsy* retrotransposons. (E) Boxplots showing 24-nt siRNA relative abundances across siren classes across cell types. Middle lines are median. Boxes span interquartile range. y-axis values are relative to per million total 24-nt siRNAs in each siRNA transcriptome. Whiskers span 2.5^th^ and 97.5^th^ percentiles. Letter grouping (α *=* 0.05) and P values are based on Tukey tests. Embryo and endosperm data from Rodrigues et al., (2013). Seedling, gametes, and pre-fertilization ovary data from Li et al., (2020).

**Fig 3:**
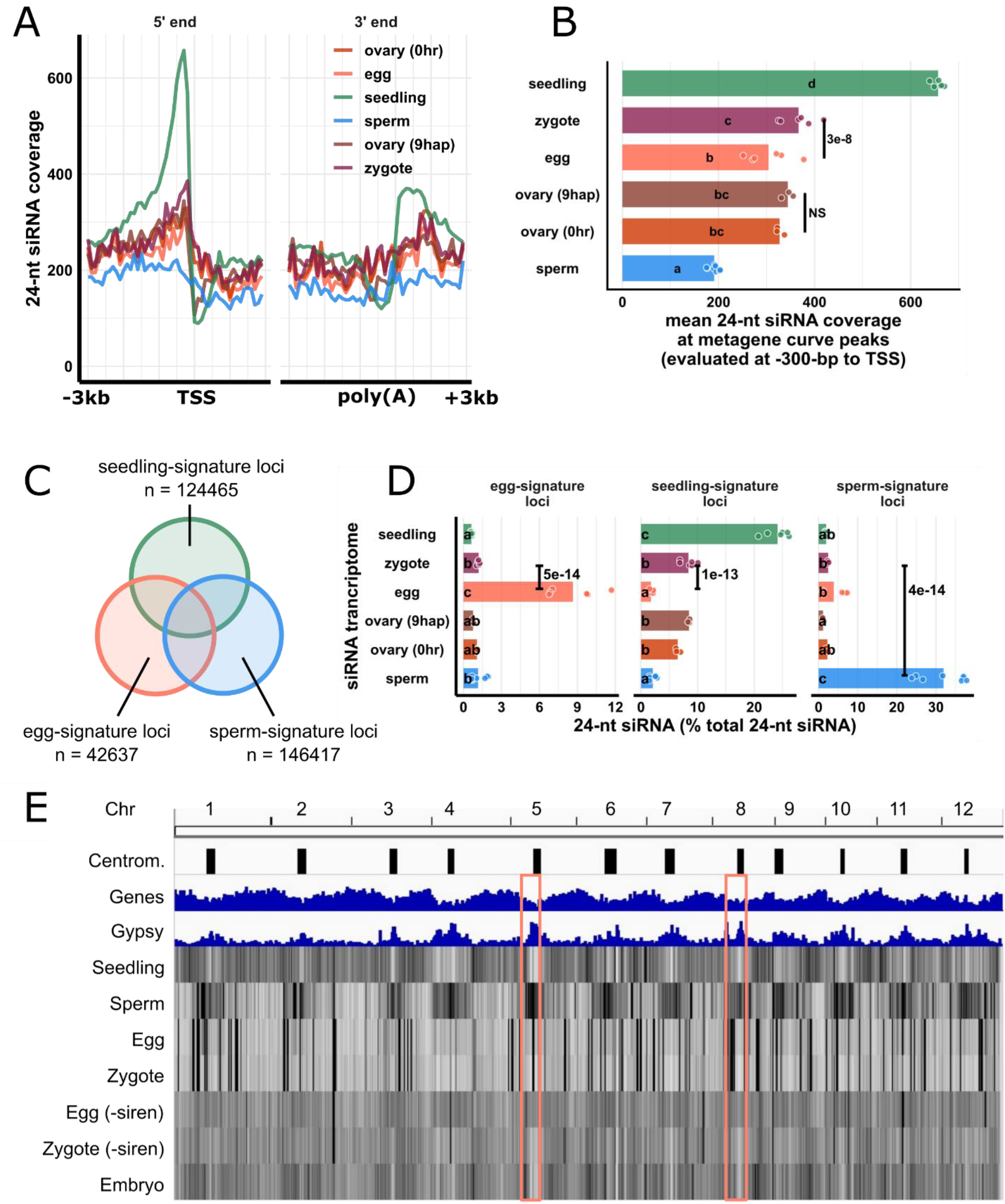
Changes in the zygote siRNA transcriptome are independent from the ovary after fertilization. (A) Metagene coverage plot for 24-nt siRNAs. Coverage is measured over 100-bp intervals and normalized per 1000 24-nt siRNAs. Vertical grid lines are 500-bp intervals. TSS transcription start site, poly(A) polyadenylation site. (B) Quantification of (A) at the interval from 300 to 200-bp upstream of TSS, corresponding to the peaks of metagene curves. Each data point is an siRNA transcriptome and bar heights are averages. x-axis values are normalized per 1000 24-nt siRNAs. (C) Venn diagram illustrating egg-signature loci (egg siRNA loci that do not overlap any seedling or sperm siRNA loci), seedling-signature loci (seedling siRNA loci that do not overlap any egg or sperm siRNA loci), and sperm-signature loci (sperm siRNA loci that do not overlap any egg or seedling siRNA loci). Sizes of overlap in Venn diagrams are not to scale. (D) Bar plot showing relative abundances of 24-nt siRNA across siRNA loci categories defined in (C). The zygote siRNA transcriptome was not used to define these locus categories. Each data point is an siRNA transcriptome. Bar heights are averages. x-axis values are normalized to total 24-nt siRNAs. (E) Heat map showing abundance of 24-nt siRNA across genome at 50-kb resolution. The first three tracks are centromeres [as defined by (Mizuno et al., 2018)], genes, and *Gypsy* retrotransposons. ‘-siren’ refers to siren siRNAs removed. Pink boxes highlight examples where egg and zygote are distinct. Letter grouping (α *=* 0.05), and P values are based on Tukey tests. Zygote and 9 hap ovary data are from this study, embryo (7-8 DAF) from Rodrigues et al. (2013), all other data from Li et al., (2020).

Next, we compared the similarity among different siren loci categories based on their genomic distributions. We used principal component analysis (PCA) to cluster the genomic distributions of the loci categories based on their abundances in 50-kb windows genome-wide (**Fig 2C**). As reference points, the genomic distributions of genes and Gypsy retrotransposons were included. On the PC plot, endosperm siren loci were well separated from all the others along PC1, which accounts for 58% of the variance in their genomic distributions. The rest of the siren loci categories were separated along PC2, which accounts for 31% of the variance, much less than what was explained by PC1. Ovary siren loci categories (pre- and post-fertilization) had similar genomic distributions, clustering closely together (**Fig 2C**). Egg and zygote siren loci also had nearly the same genomic distribution, clustering closely together (**Fig 2C**). All siren categories have distinct genomic distributions from distributions of genes or Gypsy elements (**Fig 2C**). Consistent with its unique genomic distribution, endosperm siren loci were more likely to overlap a gene (**Fig 2D**). On average, ~30% of the locus length of an endosperm siren locus was covered by a gene, whereas all the other siren categories display a similar fraction of locus length covered by genes (~13%, *P* < 1.4e-4, Tukey tests). Lastly, we compared the relative abundances of 24-nt siRNAs at different siren categories across different cell types. At endosperm siren loci, endosperm had the highest 24-nt siRNA expression, ~10-fold higher than the level in embryo and more than 100-fold higher than the levels in all other cell types we examined (**Fig 2E**). In contrast, the other siren classes shared a siRNA accumulation pattern across cell types (**Fig 2E**). Ovaries (pre- and post-fertilization), egg cell and zygote all had high abundances of 24-nt siRNAs at ovary/egg/zygote siren loci, consistent with the stable expression of egg siren siRNAs in zygote (**Fig 2B**), while seedling, sperm, embryo, and endosperm all had low abundances of 24-nt siRNAs at these siren loci. Taken together, these distinct siRNA accumulation patterns reveal that zygote siRNAs were concentrated at discrete sites similar to egg and ovary, and that the stable expression of egg siren siRNAs explains the overall similarity between zygote and egg (**Fig 1A**).

### Zygote gained siRNAs at canonical siRNA loci while losing siRNAs at egg-signature loci

Although the siRNA profile in the zygote was similar to the egg cell in terms of overall patterns of abundance, a deeper analysis revealed significant differences from the egg cell. We produced metagene siRNA coverage plots for seedling, gametes, and zygote, as well as pre- and post-fertilization ovaries (**Fig 3A**). Seedling had a strong peak upstream of the transcription start site (TSS), corresponding to where TIR transposons are enriched in the genome, with the exception of the CACTA superfamily (Han et al., 2013), and such a peak was absent in gametes and ovaries. Zygote had a significant increase in 24-nt siRNA coverage at the peak of the metagene curve relative to egg cells (**Fig 3A-B**, *P* = 3e-8, Tukey tests). In contrast, there was no significant changes between pre- and post-fertilization ovaries (**Fig 3A-B**, *P* = 0.98, Tukey tests). Thus, the differences between zygote and egg could not be due to trafficking of the newly-transcribed siRNAs from ovary. To analyze the abundance of siRNAs at individual genomic locus level, we defined siRNA loci from egg, sperm, and seedling using Shortstack (Axtell, 2013). We then classified seedling-signature loci as seedling siRNA loci that did not overlap any egg siRNA loci or sperm siRNA loci (seedling loci ∉ egg loci ∉ sperm loci, **Fig 3C**).

Overlapping siRNA loci were defined as at least 1-bp overlap in genomic coordinates (see also **Methods**). Likewise, we classified sperm-signature loci as sperm siRNA loci that did not overlap any egg or seedling siRNA loci (sperm loci ∉ egg loci ∉ seedling loci, **Fig 3C**), and lastly, egg-signature loci as egg siRNA loci that did not overlap any seedling or sperm siRNA loci (egg loci ∉ seedling loci ∉ sperm loci, **Fig 3C**). At egg-signature loci, zygote experienced a 10-fold reduction of 24-nt siRNAs (**Fig 3D**, *P* = 5e-14, Tukey tests). At seedling-signature loci, zygote had 4.7-fold more 24-nt siRNAs than egg cell (**Fig 3D**, *P* = 1e-13, Tukey tests). Gaining siRNAs at gene-proximal regions and seedling-signature loci is consistent with an increase of TIR siRNAs in zygote (**Fig 1C**). Since these seedling-signature loci did not overlap any egg siRNA loci or sperm siRNA loci, the increase of 24-nt siRNAs at seedling-signature loci in zygote was unlikely due to carryover from either gamete. At sperm-signature loci, zygote had very few 24-nt siRNAs (**Fig 3D**, zygote vs. sperm *P* = 4e-14, Tukey tests), much like the results for miRNAs (**Fig 1E**), confirming small RNA contribution from sperm cell is very limited relative to egg. There was little difference in the ovaries before and after fertilization for any of these locus categories (**Fig 3D**, *P* = 0.76, *P* = 0.84 and *P* = 0.84 at egg-, seedling- and sperm-signature loci, respectively). It is important to note that the zygote siRNA transcriptome was not used to define these locus categories. Lastly, we bioinformatically removed siren siRNAs from egg and zygote libraries (**Fig 3E**) and re-analyzed their genome-wide 24-nt siRNA distributions. This analysis revealed that outside of the siren loci (which as defined previously constitute ~1% of all 24-nt siRNA loci), zygote and egg were indeed distinct from each other in genome-wide 24-nt siRNA distribution (**Fig 3E** and **Fig S2F**, *P* = 0, Tukey Tests). Egg cell has a slight enrichment of 24-nt siRNAs at centromeric regions, while zygote showed a relative depletion of siRNAs at centromeric regions (**Fig S2G**, egg vs. zygote *P* = 0, Tukey tests), much like embryo (zygote vs embryo *P* = 0.5, Tukey tests; **Fig 3E**, pink boxes indicate two examples, see also **Fig S2G**). Taken together, these results indicate that the zygote has an siRNA transcriptome that is distinct from that of the egg cell, and further, that the changes from egg cell to zygote were independent of post-fertilization changes in the ovary.

To further characterize the differences between the zygote siRNA transcriptome and that of the egg cell, we next defined zygote siRNA loci using Shortstack with zygote siRNAs. We then classified Z-E loci as zygote siRNA loci that did not overlap any egg cell siRNA loci (Z loci ∉ E loci in set operation), E-Z loci as egg siRNA loci that did not overlap any zygote siRNA loci (E loci ∉ Z loci), and Z/E loci intersect as zygote siRNA loci that overlapped egg siRNA loci (Z loci ∩ E loci, **Fig 4A**). Despite the similarities between egg and zygote at the high abundance siRNA loci (**Fig 1A**, **Fig 2B**), widespread distinct siRNA loci were detected in one cell type but not the other. There were 101,841 E-Z loci (newly diminished siRNA loci in zygote), 94,591 Z-E loci (newly detected siRNA loci in zygote), but only 42,437 Z/E loci intersect.

**Fig 4:**
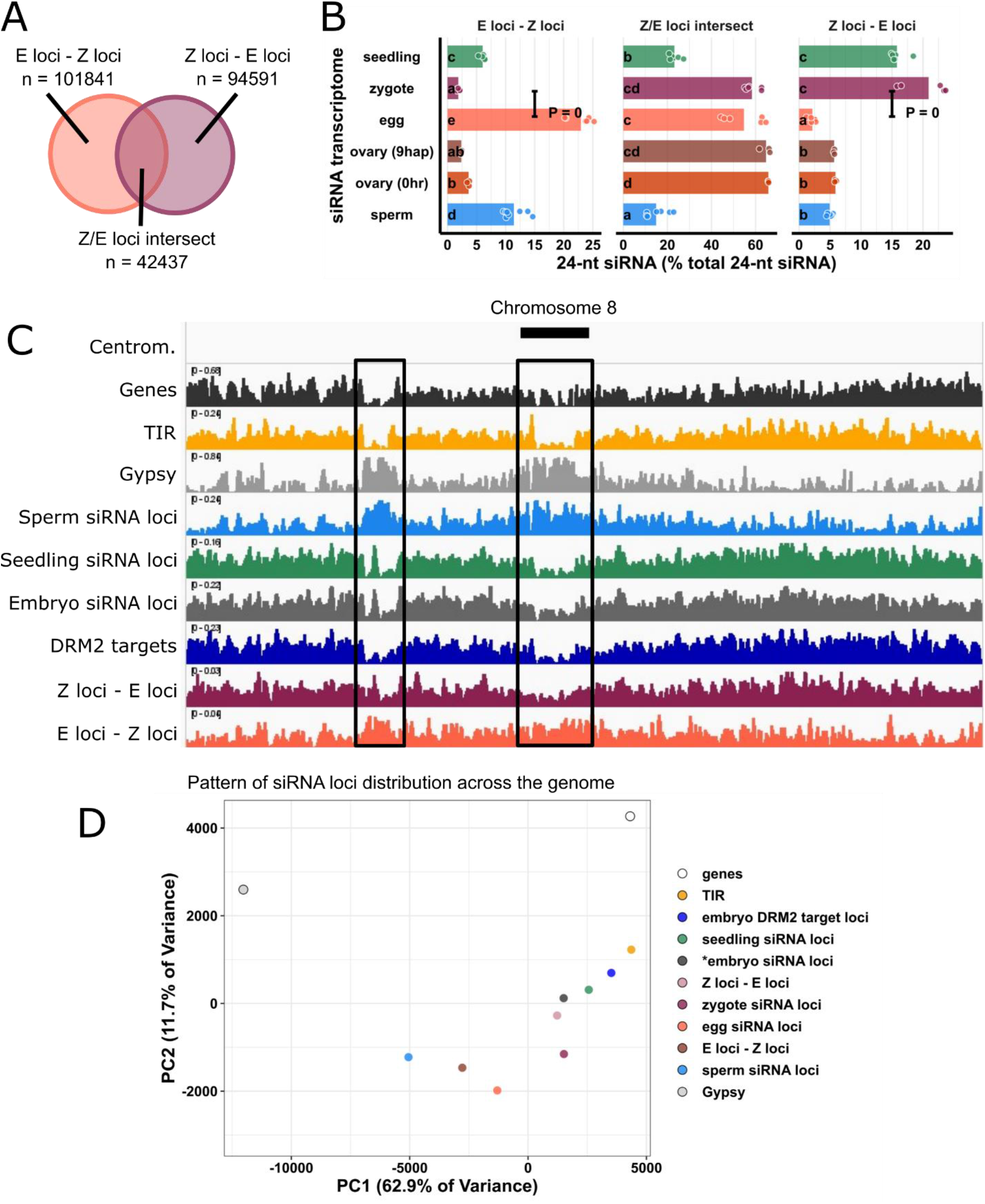
Widespread newly detected siRNA loci in zygote. (A) Venn diagram illustrating E-Z loci (egg siRNA loci that do not overlap any zygote siRNA loci, E loci ∉ Z loci), Z-E loci (zygote siRNA loci that do not overlap any egg siRNA loci Z loci ∉ E loci), and Z/E loci intersect (zygote siRNA loci that overlap egg siRNA loci, Z loci ∩ E loci). Sizes of overlap in Venn diagrams are not to scale. (B) Quantification of 24-nt siRNA relative abundances for (A). Each data point is a siRNA transcriptome. Bar heights are averages. x-axis-values are relative to total 24-nt siRNA reads. Letter grouping (α = 0.05), and P values are based on Tukey tests. (C) Distribution of siRNA loci along a chromosome. Chromosome 8 is chosen because it is one of the chromosomes with a completed sequenced centromeric region (Mizuno et al., 2018). Centrom. Centromeric regions; TIR: terminal inverted repeat transposons, CACTA superfamily excluded. *Gypsy*: *Gypsy* retrotransposons. Black boxes highlight regions with abundant *Gypsy* retrotransposons and relative depletion of TIR, seedling siRNA loci, embryo siRNA loci, DRM2 targets, and Z-E loci. (D) Principal component plot showing siRNA loci distribution across the genome. Distributions are evaluated at 50-kb resolution across the genome. Each data point is the distribution of a loci category. Zygote and 9 hap ovary data are from this study, all other data from Li et al., (2020).

When 24-nt siRNA reads at individual loci were tallied and normalized to total 24-nt siRNAs, as expected, we found that at E-Z loci, egg had ~10-fold more 24-nt siRNAs than the zygote (**Fig 4B**, *P* = 0, Tukey tests); at Z-E loci, zygote had ~10-fold more 24-nt siRNAs than egg (**Fig 4B**, *P* = 0, Tukey tests); and no difference at Z/E loci intersect. There were siRNAs not captured by siRNA loci. These siRNAs resided at genomic regions with insufficient siRNAs and did not meet the 0.5 RPM threshold for assignment as loci on Shortstack (see also **Methods**), which explains the small number of egg siRNAs at Z-E loci and the small number of zygote siRNAs at E-Z loci. There were no differences between ovaries before and after fertilization in any of the three locus categories (**Fig 4B**), again suggesting changes in the zygote siRNA transcriptome were not coupled with the ovary (**Fig 3**). In addition, the abundance of seedling siRNAs in Z-E loci and scarcity in E-Z loci revealed the emergence of a seedling-like siRNA pattern in zygote (**Fig 4B**). Since the seedling siRNA transcriptome was not used to classify Z-E loci, this observation suggests that the zygote has initiated a return to the canonical siRNA profile, consistent with the increase in 24-nt siRNAs from TIR transposons (**Fig 1C**) as well as at gene-proximal regions (**Fig 3A-B**) and seedling-signature loci in zygote (**Fig 3D**).

During ZGA of mRNA transcriptomes, genes expressed in zygote but not in egg cell all had initially low expression relative to a background of abundant maternal transcript carryover (Anderson et al., 2017; Chen et al., 2017; Zhao et al., 2019). Thus, if the siRNA transcriptome transitions similarly in the zygote, one would expect to see an initial widespread detection of low abundance 24-nt siRNAs at new loci, relative to a background of more abundant maternal carryover siRNAs corresponding to egg siren loci. Indeed, in contrast to the high abundance siRNAs of intersect loci, Z-E loci and E-Z loci overall had lower siRNA abundances than zygote/egg intersect loci (**Fig 4B**). Nevertheless, on average, one in every five zygote 24-nt siRNAs (~20%) resided at Z-E loci in the zygote. Together with the numerical abundance of Z-E loci (70% of all zygote loci) these results suggest that newly detected siRNA loci in zygote are widespread and explain a substantial fraction of 24-nt siRNAs in zygote.

The highly expressed siren loci in egg and zygote raise the concern of whether the apparent upregulation of Z-E loci could be explained by downregulation of egg siren loci. Consistent with the stable expression of egg siren loci in zygote, including or excluding siren siRNAs did not change the results of analyses (**Fig S2**, **Fig 3A-D**, **Fig 4A-B**), supporting the distinct distributions of non-egg-siren 24-nt siRNAs in egg and zygote (**Fig 3E**). Taken together, changes in the zygote siRNA transcriptome are not explained by downregulation of abundance egg siren siRNAs, but due to up- and downregulation of other siRNA loci that are widespread across genome.

### Newly-detected siRNA loci in zygote resemble canonical siRNA loci in terms of genomic location and DNA methylation

To investigate the patterns and characteristics of zygote siRNA loci, we compared the genomic distribution of zygote siRNA loci and Z-E loci against a set of different siRNA loci categories, including E-Z loci, egg siRNA loci, embryo siRNA loci (Rodrigues et al., 2013), seedling siRNA loci, and sperm siRNA loci. Our efforts to generate robust DNA methylome profiles for zygotes were not successful, possibly because zygotes are fragile as compared to egg cells, and the output of random-primed based methylome sequencing methods are highly sensitive to library preparation conditions (Li et al., 2019). However, we had previously generated a rice *drm2* mutant using CRISPR-Cas9 genome editing. By comparing mCHH between mature wildtype and *drm2* embryos, we had previously identified a set of DRM2 targets (Li et al., 2020), which define canonical sites of RdDM. As reference points, we also included genes, TIR transposons and *Gypsy* retrotransposons (Kawahara et al., 2013). Z-E loci resembles the distribution of TIR, seedling and embryo siRNA loci, as well as embryo DRM2 targets. In contrast, the E-Z loci representing the newly diminished loci in zygote had a distinct pattern, more similar to sperm siRNA loci, which is more heterochromatic (**Fig 4C**). There was a relative depletion of siRNA loci from centromeric regions for zygote siRNA loci, much like canonical siRNA loci, and unlike egg siRNA loci (**Fig S3A**, *P* = 4e-10, Tukey tests). Consistent with a more similar distribution to canonical siRNA loci, zygote siRNA loci and Z-E loci had higher degrees of overlap with seedling siRNA loci and embryo DRM2 targets, while egg siRNA and E-Z loci had low degrees of overlap, as did sperm siRNA loci (**Fig S3B**). Zygote siRNA loci and Z-E loci overlapped larger numbers of DRM2 targets per Mb genome space, much like seedling siRNA loci, and unlike egg siRNA loci, E-Z loci or sperm siRNA loci (**Fig S3C**).

To gain more information on the factors underlying the variation in siRNA loci distributions, we used principal component analysis (PCA) to cluster the genomic distributions of the above locus categories based on their abundance in 50-kb windows genome-wide (**Fig 4D**). The separation of locus categories along PC1 had a near-perfect rank order correlation with median distance to nearest genes (**Fig S3D**, rho = −0.98, *P* = 0), and PC2 was correlated with median length of locus categories (**Fig S3E**, rho = 0.88, *P* = 7e-4). PC1, which explained 63% of variance in genomic distributions across loci categories, was strongly correlated with various aspects of rice genome organization. PC1 was strongly correlated with TIR transposon overlap as well as mCHH level in wildtype embryo (**Fig S4A** and **C**). PC1 was also strongly anti-correlated with *Gypsy* retrotransposon overlap and mCG, and to a lesser extent mCHG in wildtype embryo (**Fig S4B, D** and **E**). These genomic features are mutually correlated (**Fig S4F**), consistent with the prior understanding of cereal genome organization (Gent et al., 2013; Han et al., 2013).

Strong correlations between PC1 (gene proximity), TE overlap, and DNA methylation led us to statistically assess the differences of these attributes among siRNA loci categories. TIR transposons, where RdDM is known to take place in cereal genomes, was gene proximal, consistent with the gene proximal distribution of embryo DRM2 targets (**Fig 5A**). Canonical siRNA loci, such as seedling siRNA loci and embryo siRNA loci, were closer to genes than non-canonical siRNA loci, such as sperm siRNA loci (**Fig 5A**). E-Z loci, the newly diminished siRNA loci in zygote, were on average much farther away from genes than Z-E loci were (**Fig 5A**, 2.8-kb vs. 1.6-kb, *P* = 0 Tukey tests), consistent with their heterochromatic genomic distributions (**Fig 4C-D**, **Fig S3A**). Total zygote siRNA loci were closer to genes than total egg siRNA loci (*P* = 0). From egg cell to zygote, there was a 30% decrease in median distance (2.4-kb vs. 1.6-kb). In contrast, from zygote to embryo (7 days after fertilization, data from Rodrigues et al., 2013), there was an 6% decrease (1.6-kb vs. 1.5-kb). In gametes, siRNA loci were more likely to overlap a *Gypsy* retrotransposon than a TIR transposon (**Fig 5B**). However, in sporophytes, including zygote itself, siRNA loci are more likely to overlap a TIR transposon than a *Gypsy* retrotransposon (**Fig 5B**). These observations are consistent with the results where zygote had more TIR siRNAs and less *Gypsy* siRNAs (**Fig 1C**), more gene-proximal 24-nt siRNAs than egg cell (**Fig 3A-B**), zygote had increased siRNAs in seedling-signature loci (**Fig 3C**), and seedling had comparable siRNA level with zygote in Z-E loci (**Fig 4B**).

**Fig 5:**
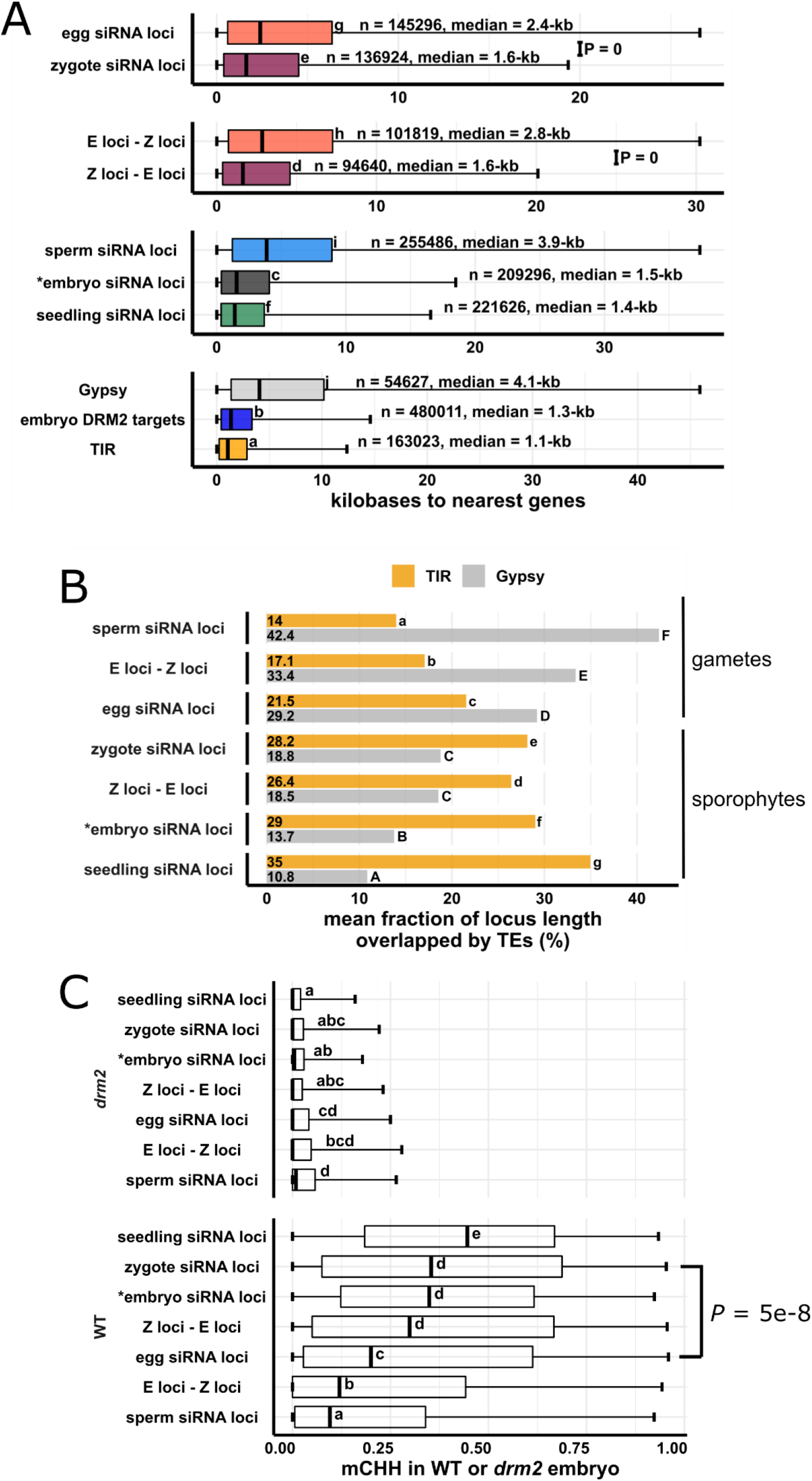
Newly detected siRNA loci in zygote reset to the canonical siRNA profile and predict CHH methylation in embryo in an RdDM-dependent manner. (A) Boxplots showing distance of siRNA loci to nearest genes. Middle lines are median. Boxes span interquartile range. Whiskers span 2.5^th^ and 97.5^th^ percentiles. (B) Bar plots showing mean locus length overlapped by TIR or *Gypsy* transposons across siRNA loci categories. Statistical comparisons are made across siRNA loci categories within a TE superfamily. (C) Boxplots showing CHH methylation level in mature wildtype and *drm2* mutant embryos. Middle lines are median. Boxes span interquartile range. Whiskers span 2.5^th^ and 97.5^th^ percentiles. E-Z loci: *n* = 101,841, Z-E loci: *n* = 94,591 (69% of all zygote siRNA loci). Letter groupings (α = 0.05) and P values are based on Tukey tests. *Embryo siRNA data from Rodrigues et al., (2013), which was based on a single replicate. Seedling, gametes, and pre-fertilization ovary data from Li et al., (2020).

Resetting to the canonical siRNA pattern might suggest that the newly detected siRNA loci in the zygote are targeted for CHH methylation during embryogenesis in an RdDM-dependent manner. We compared DNA methylation levels across different siRNA loci categories in mature wildtype and *drm2* embryos (**Fig 5C**, see also **Fig S5**). Although all siRNA loci categories were associated with RdDM during embryogenesis, as median mCHH levels were all higher in wildtype embryo than *drm2* embryo, zygote siRNA loci had much higher level of mCHH than egg siRNA loci did in wildtype embryo (**Fig 5C**, *P* = 5e-8), much like embryo and seedling siRNA loci. Importantly, newly detected siRNA loci (as represented by Z-E loci) had high mCHH levels in wildtype embryo (*P* = 0), whereas newly diminished siRNA loci (E-Z loci) had low mCHH levels in wildtype embryo. In addition, while there were smaller differences in mCHG and especially in mCG across locus categories in wildtype embryo when compared to mCHH, the full extent of methylation across siRNA loci categories also depended on DRM2 for all three contexts (**Fig S5**). It is important to note that zygote siRNA loci had high degrees of overlap with canonical RdDM loci (seedling siRNA loci and embryo DRM2 targets, **Fig S3B**), and that zygote siRNA loci overlapped large number of DRM2 targets per Mb genome space (**Fig S3C**). Together with elevated mCHH level in embryo, these results suggest that newly detected zygote siRNA loci mark canonical siRNA loci that will undergo hypermethylation during embryogenesis, rather than defining hypermethylated regions that are distinct from canonical siRNA loci.

Lastly, the fact that siren loci were defined by abundant 24-nt siRNAs (**Fig 2A**) led us to speculate that they would have high mCHH. In fact, we found the opposite, i.e., they had lower mCHH levels relative to other siRNA loci in each tissue (**Fig S6**). In ovary, siren loci had lower mCHH level than canonical siRNA loci (seedling siRNA loci), and ovary siRNA loci that were not siren loci had comparable mCHH level to siren loci, if not higher (**Fig S6A**). Similar results were found for egg cell and embryo as well (**Fig S6B-C**). At siren loci as well as siRNA loci that were not siren loci, DMR2 was required for mCHH in embryo. Lastly, both wildtype and *drm2* endosperm had overall low mCHH methylation, and endosperm siren loci did not correspond to high mCHH level in the endosperm (**Fig S6D**). Unlike mCHH, mCG and mCHG did not produce any notable pattern across cell types (**Fig S6**). Taken together, these results suggest that although the highly abundant siRNAs produced by siren loci may also target DNA methylation in a DRM2-dependent manner, they do so inefficiently as compared to siRNAs from canonical RdDM loci.

## Discussion

### The parental gametes have unequal contributions to the zygote siRNA transcriptome in rice

The zygote is a critical stage in the transition from gametophyte to sporophyte, but despite long standing evidence for overall chromatin reorganization after fertilization (Ingouff et al. 2010) there is little information available on the epigenetic changes in the zygotic genome that accompany this transition. We report here the first small RNA transcriptome characterization of plant zygotes. Overall, the small RNA transcriptome of the zygote is similar to that of the egg cell (**Fig 1A-D**), which we explain by transcript carryover from the egg cell, and the dilution of sperm cell small RNAs. This is supported by our finding that the most abundant miRNAs in egg continued to be the most abundant in zygote (**Fig 1D**). In contrast, top sperm-enriched miRNAs became orders of magnitudes less abundant in zygote (**Fig 1E** and **Fig S1C**), and zygote had very few 24-nt siRNAs at sperm-signature siRNA loci **(Fig 3D**). These observations have ensuing implications for the current models of post-fertilization silencing through the male germline. It has been proposed that sperm-transmitted siRNAs regulate TEs and balance parental contribution in the endosperm, as RdDM-mutants affect endosperms from 2n maternal × 4n paternal crosses (Borges et al., 2018; Erdmann et al., 2017; Martinez et al., 2018; Satyaki and Gehring, 2019) in *Arabidopsis*. Our data indicate that at least in rice embryogenesis, any effects of sperm-transmitted siRNAs on embryos are likely to be indirect. Assuming sperm-derived siRNAs are also diluted by the larger central cell, we speculate that the effect of the sperm-transmitted siRNAs may act through sperm chromatin modifications and not siRNAs themselves. In the newly formed endosperm, there is lack of an active replacement of histone variants, and sperm-derived histone variants are passively diluted through nuclear divisions (Ingouff et al., 2007, 2010). In contrast, in the zygote, histone variants are actively replaced in a replication-independent manner before the first embryonic division (Ingouff et al., 2007, 2010).

### A special class of highly abundant maternal siRNAs persists in the zygote

We found that a small number of loci accounted for most of the siRNAs in egg cells and zygotes (**Fig 2A**). We refer to these loci as siren loci, using the term applied to similar loci in rice endosperm (Rodrigues et al., 2013), and more recently in *Brassica rapa* and *Arabidopsis* ovules (Grover et al., 2020). Importantly, egg siren siRNAs remained similarly highly expressed in the zygote (**Fig 2B**). Thus, upregulation of siRNAs at newly detected zygote siRNA loci cannot be explained by large downregulation of egg siren loci, which was further demonstrated by similar results for these newly detected loci obtained after excluding the siren siRNAs from the analysis (**Fig S2**). Siren loci were first discovered in rice endosperm (Rodrigues et al., 2013). In *Arabidopsis* and *Brassica rapa* (Grover et al., 2020), siren loci detected in ovules are also highly expressed in the endosperm; however, siren loci in rice ovary are distinct from those detected in rice endosperm (**Fig 2B-E**). Siren loci in the zygote were distinct from endosperm siren loci in endosperm collected 7-8 days after fertilization, instead coinciding with siren loci detected in ovary and egg cell (**Fig 2B-E**). However, it remains possible that the central cell and earlier stages of endosperm have an siRNA transcriptome more like that of the zygote. It has been proposed that the embryo receives siRNAs from the endosperm (Hsieh et al., 2009; Martínez and Köhler, 2017). This does not appear to be the case in 7-8 day rice seeds, since rice embryos had low siRNA abundance at endosperm siren loci at this stage (Rodrigues et al., 2013). A recent publication demonstrated that trans-acting siRNAs from ARFs (tasiR-ARF) traffic across ovule cell layers to regulate megaspore mother cell (MMC) identity in *Arabidopsis* (Su et al., 2020). It has also been proposed that siRNAs may traffic from the seed coat into the embryo during seed development (Grover et al., 2020, 2018). Likewise, it is possible that siren siRNAs in the egg cell and zygotes are produced in the ovary tissue instead. Although siRNAs at siren loci may direct some CHH methylation in ovary or during embryogenesis, they appeared to have lower mCHH level than their non-siren siRNA loci counterparts (**Fig S6**). Therefore, it is unlikely that siren siRNAs play a role in embryogenesis through directing DNA methylation. However, we cannot exclude the possibility that the 24-nt siRNAs of siren loci function in chromatin modification or post transcriptional silencing independently of DNA methylation in the zygote or egg, regardless of their cell type of origin.

### The siRNA transcriptome landscape of rice zygotes indicates resetting towards a canonical siRNA pattern

We detected widespread new zygote siRNA loci relative to the egg cell, representing ~69% of all zygote siRNA loci. There were 94,591 zygote siRNA loci that did not overlap any egg siRNA loci (Z loci ∉ E loci or Z-E loci, **Fig 4A**), as compared to 42,437 siRNA loci that overlapped egg siRNA loci (Z loci ∩ E loci or Z/E loci intersect, **Fig 4A**). In addition, 101,841 egg siRNA loci were diminished in zygote (E loci ∉ Z loci or E-Z loci, **Fig 4A**). In relative abundance, most of the siRNA reads were accounted for by egg siRNA carryover and stably-expressed egg siren siRNAs (**Fig 2B**), and thus siRNA abundance was lower at Z-E loci than at Z/E loci intersect which contains the siren siRNAs (**Fig 4B**). This low relative abundance can be understood in the context of the zygotic transition, which involves a new genomic program initiated within that one cell, so that production of new siRNAs, either by RNA polymerase IV or RNA polymerase II, will be occurring against the backdrop of egg cell RNA carryover. Similar observations have been made for zygote mRNA transcriptomes from multiple independent laboratories from different plant species (Anderson et al., 2017; Chen et al., 2017; Zhao et al., 2019), where zygote *de novo* expressed genes, including those with key functions in embryogenesis, were lowly expressed in the zygote. In fact, the relative abundance of the zygote siRNAs at Z-E loci (~20%) is similar to that of egg cell siRNAs at E-Z loci (~22.5%; **Fig 4B**), but they differ significantly in their genome-wide distribution as discussed below.

Several lines of evidence indicate the zygote has initiated a resetting towards the canonical siRNA pattern, and that such resetting is independent from the ovary. First, the zygote had increased 24-nt siRNA from TIR transposons, and decreased siRNAs from *Gypsy* retrotransposons, as compared to the egg cell (**Fig 1C**). Second, zygote had increased 24-nt siRNAs at seedling-signature loci (**Fig 3D**), as compared to the egg cell. In contrast, there were no significant changes to the 24-nt siRNAs at seedling-signature loci in ovaries pre- and post-fertilization (**Fig 3D**). Moreover, seedling had comparable 24-nt siRNAs to zygote at Z-E loci (**Fig 4B**). As the zygote siRNA transcriptome was not used to define seedling-signature loci, and the seedling siRNA transcriptome was not used to define Z-E loci, these results serve as an objective indication that the zygote shifted towards a more seedling-like siRNA transcriptome. Third, zygote had increases relative to the egg cell in 24-nt siRNAs at the TSS region upstream of genes, while there was lack of a corresponding change in the ovary after fertilization (**Fig 3A-B**). High 24-nt siRNA coverage upstream of genes around the TSS is a feature of a canonical siRNA transcriptome, as exemplified by seedling (**Fig 3A**). Fourth, the genomic distribution of Z-E loci is more similar to TIR transposons, embryo siRNA loci, embryo DRM2 targets and seedling siRNA loci, while that of E-Z loci is not (**Fig 4C-D**). Consistent with the major contribution of Z-E loci to the zygote siRNA distribution, the total set of zygote siRNA loci also displayed a closer relationship to the canonical siRNA distribution than did the total egg siRNA loci (**Fig 4C**). Fifth, similarities in genomic distribution were confirmed by distances to the nearest genes and TE overlaps (**Fig 5A-B**). Notably, there was a 30% decrease in median distance to genes from egg to zygote, which took place over the course of less than one cell cycle. Only an 6% decrease occurred during the transition from zygote to embryo (7 days after fertilization), occurred over the course of numerous cell cycles. Consistent with distance to nearest genes, gamete siRNA loci were more likely to overlap a *Gypsy* element than a TIR element; while zygote siRNA were more likely to overlap a TIR element instead, like the rest of sporophyte siRNA loci categories.

### Newly detected zygote siRNAs mark future CHH hypermethylation sites in mature embryos

Hypermethylation of embryo has been reported in a number of angiosperm species, including *Arabidopsis*, soybean, chickpea, *Brassica rapa*, and rice (Bouyer et al., 2017; Chakraborty et al., 2020; Kawakatsu et al., 2017; Li et al., 2020b; Lin et al., 2017; Rajkumar et al., 2020; Zhou et al., 2021). We found that newly detected siRNA loci have abundant CHH methylation in embryos that is dependent on the RdDM methyltransferase DRM2 (**Fig 5C**). Although all siRNA loci categories had higher mCHH levels in wildtype embryo than in *drm2* embryo, zygote siRNA loci had higher mCHH levels than egg siRNA loci in mature wildtype embryo, resembling embryo and seedling siRNA loci. These results indicate that newly detected siRNA loci in zygote not only reset to canonical siRNA pattern, but also that the corresponding 24-nt siRNAs are capable of targeting high CHH methylation during embryogenesis. Since zygote siRNA loci have a similar distribution to canonical siRNA loci (**Fig 4D**) and had substantial degrees of overlap with seedling siRNA loci and embryo DRM2 targets (**Fig S3B-C**), zygote siRNAs are associated with high CHH methylation at regions similar to seedling siRNA loci and embryo DRM2 targets, instead of regions independent from canonical siRNA loci. Thus, resetting of the gametic 24nt siRNA loci to a distribution that results in embryo hypermethylation appears to be initiated in the zygote before the first embryonic division. Reminiscent of the increased heterochromatic siRNAs in rice gametes, a recent paper in *Arabidopsis* revealed that heterochromatin is decondensed during embryogenesis and promotes a transient production of siRNAs from heterochromatic TEs at least as early as the preglobular stage (Papareddy et al., 2020). The *Arabidopsis* embryo siRNAs from euchromatic TE and canonical siRNA loci peaked towards the end of embryo morphogenesis. However, as neither egg nor zygote siRNAs have been sequenced yet in *Arabidopsis*, the relationship of the heterochromatic siRNAs in *Arabidopsis* embryos to the siRNA reprogramming in egg cells and zygotes that is described here for rice remains to be determined.

## Conclusions

Plant gametes are highly dimorphic in terms of size, chromatin (Wang and Köhler 2017; Borg and Berger 2015; Ingouff et al. 2010), and gene expression (Anderson et al., 2013), consistent with a differential reprogramming of gamete epigenomes prior to fertilization inferred from their siRNA profiles (Li et al., 2020). In mammals, studies have found a progressive change in epigenomes after the two-cell embryo stage and concluded by the blastocyst stage (Xu and Xie, 2018). Due to the extreme difficulties associated with plant zygote isolation and corresponding low-input sequencing, epigenome profiles of zygotes have remained poorly characterized in plants. The results from this study in rice suggest that while the zygote inherits maternal (but not paternal) siRNAs, the resetting to the canonical siRNA transcriptome is initiated, setting the stage for the methylation pattern in the embryo. This conclusion is consistent with previous observations in *Arabidopsis* that replacement of H3 variants occurs in the zygote before the first cell division (Ingouff et al., 2007; 2010). Lastly, as siRNA expression is influenced by histone modifications, and siRNAs can either reinforce or initiate DNA methylation and histone modifications, the siRNA transcriptome is an indicator and output of the epigenome. Thus, it appears likely that resetting of the other features of the epigenome, such as histone modifications and chromatin conformation, may also be initiated in plant zygotes after fertilization.

## Methods

### Plant growth condition and zygote collection

Rice (*Oryza sativa*) variety *Kitaake* was grown in soil in greenhouse under natural light condition. Zygote isolation was performed as described (Anderson et al., 2017; Li et al., 2019). Briefly, rice flowers were hand pollinated. At eight to nine hours post pollination, ovaries were dissected. A transverse cut was made at the middle region of the ovary in a droplet of 0.3 M mannitol. The lower part of the cut ovary was gently pushed using an acupuncture needle to separate selected cells under a phase contrast inverted microscope. Once the zygote was separated and floated out of the ovary incision, it was captured by a fine glass capillary and immediately frozen in liquid nitrogen. We routinely culled any unfertilized egg cells that did not conform to zygotic cell morphology during our collections (Anderson et al., 2017). 50 zygotes were collected for each replicate, and six replicates were collected. Intact ovaries at 8-9 hours after pollination were collected separately for the ovary small RNA analysis. 10 ovaries were collected for each replicate, and three replicates were collected (**Supplemental Table 1**).

### RNA extraction and small RNA library construction

RNA extractions were performed using Ambion RNAqueous Total RNA kit (AM1931), including an on-column DNase I treatment using Qiagen DNase I (79254). Total RNA was analyzed using a Bioanalyzer (Agilent) to check for RNA integrity, with the eukaryotic total RNA-pico program. RNA input for library construction was ~30 ng. Small RNA libraries were made using the NEXTflex small RNA-seq kit v3 (PerkinElmer NOVA-5132-05), with the following modifications. ¼ dilution of adapters was used. The 3’ adapter ligation step was done at 20°C overnight. Zygote libraries were amplified at 24 cycles. Post-fertilization ovary libraries were amplified at 20 cycles, as pre-fertilization ovaries (Li et al., 2020). The library product was size selected using PippinHT (Sage Science) 3% agarose gel cassettes.

### Small RNA sequencing analysis

Analyses were based on the Os-Nipponbare-Reference-IRGSP-1.0 reference genome (Kawahara et al., 2013). Genome annotations for transposable elements, genes, miRNAs, 5S rRNA, tRNA, NOR, CentO repeats and phasiRNA loci were performed as described (Li et al. 2020). Quality filtering, adapter trimming, PCR duplicate removal and alignment were performed as described (Li et al. 2020). Small RNA-seq reads were quality filtered and trimmed of adapters using cutadapt (Martin, 2011), parameters “-q 20 -a TGGAATTCTCGGGTGCCAAGG -e .05 -O 5 --discard-untrimmed -m 28 -M 33”. PCR duplicates were then removed using PRINSEQ, parameters “prinseq-lite.pl -fastq out_format 3 - out_good -derep 1” (Schmieder and Edwards, 2011). The four random nucleotides at each end were then removed using cutadapt “-u4” followed by cutadapt “-u −4”. Reads were aligned to the genome with BWA-backtrack (version 0.7.15) (Li and Durbin 2009), parameters “aln -t 8 −l 10.” Except where indicated otherwise, multi-mapping reads were included in all analyses. The uniquely mapping subset of siRNAs was defined by having MAPQ values of at least 20 using SAMtools (Li et al. 2009). Except where indicated otherwise, siRNAs used for analyses were small RNA reads (20 – 25-nt) not overlapping 90% or more of their lengths with miRNA, 5S rRNA, tRNA, NOR and phasiRNA loci as determined by the BEDTools coverage tool (Quinlan and Hall, 2010). For analysis of overlaps of siRNAs at *Gypsy* retrotransposons, the CentO centromeric tandem repeat, Terminal Inverted Repeat (TIR) DNA transposons, and 24-nt siRNA loci, only siRNAs that overlapped by at least 50% of their lengths were counted. CACTA elements were excluded from the TIR DNA transposons. Distances to closest genes were obtained using the BEDTools closest tool. Whole-genome small RNA heat maps were made on 50-kb intervals using IGVtools (Thorvaldsdottir et al., 2013). For better visualization of midrange values, heatmap intensity was maxed out at 1.25× coverage per 10 million 24-nt siRNAs.

### miRNA analysis

To measure miRNA accumulation, the BEDTools coverage tool was used to count the number of 20 – 25-nt reads that overlapped at least 90% of their length with annotated miRNA positions (**Supplemental Dataset 1)**. R package EdgeR was used to analyze miRNA accumulation (McCarthy et al., 2012). Individual miRNA counts were normalized by total mapped small RNAs and filtered for >1 counts per million reads (CPM) in at least three libraries. Differential expression analyses were performed under |log2FC| > 1 and FDR < 0.05 cutoffs. Differential expressing miRNA genes were visualized under counts per million miRNAs.

### Definition of siRNA loci

Small RNA loci were identified from the initial 20 – 25-nt total small RNA alignment BAM files using Shortstack (Axtell, 2013) after merging replicates using default parameters. Each cell type was downsampled to 3.5 million small RNAs first. For each tissue type (pre- and post-fertilization ovary, egg cell, sperm cell, zygote, seedling, embryo and endosperm), siRNA loci were defined as RPM > 0.5, 24-nt-dominant and not detected as a miRNA locus (‘DicerCall=24; MIRNA=N’). Endosperm siren loci were defined as the highest expressing loci that accounted for 60% of the cumulative RPM in the endosperm. Similarly, pre- and post-fertilization ovary siren loci as well as egg and zygote siren loci were defined as the highest expressing loci that accounted for 60% of the cumulative RPM in the ovary. The 60% cutoff was selected based on the turning point of cumulative expression vs. percentage rank plot of ovary (**Fig 2A**). Seedling-signature loci were identified as seedling siRNA loci that did not overlap any sperm siRNA loci or egg siRNA loci (seedling loci ∉ egg loci ∉ sperm loci, **Fig 3C**) using the BEDTools intersect tool (Quinlan and Hall, 2010). Overlaps were defined as at least 1-bp overlapping genomic coordinates. Similarly, sperm-signature loci were identified as sperm siRNA loci that did not overlap any egg siRNA loci or sperm siRNA loci (sperm loci ∉ egg loci ∉ seedling loci, **Fig 3C**). Egg-signature loci were identified as egg siRNA loci that did not overlap any seedling siRNA loci or sperm siRNA loci (egg loci ∉ seedling siRNA loci ∉ sperm siRNA loci, **Fig 3C**). Z-E loci were zygote siRNA loci that did not overlap egg siRNA loci (Z loci ∉ E loci). E-Z loci were egg siRNA loci that did not overlap zygote siRNA loci (E loci ∉ Z loci). Z/E loci intersect were zygote siRNA loci that overlapped egg siRNA loci (Z loci ⋂ E loci, **Fig 4A**).

### DNA methylation analyses

Methylation values were calculated for each locus using the mtr function of CGmapTools v0.1.2 (Guo et al., 2018) using the CGmap files generated in our previous study as input (Li et al., 2020). Only loci with more than 3 (mC + C) calls were included in the analyses.

### Statistical analyses

Tukey tests were performed using the R package emmeans (Searle et al., 1980) with multiple comparison correction using Tukey’s method. Letter groupings were done at α = 0.05, where the differences between means sharing the same letter were not statistically significant. For multifactorial analyses, multiple comparisons were applied to families of tests at each interacting factor level: at the level of each TE/locus category for **Fig 1C**, **Fig 2E**, **Fig 3D**, **Fig 4B**, **Fig 5B**, **Fig S1A**, **Fig S2D**, **Fig S2F**, and **Fig S3B**, and at the level of genotype and context for **Fig 5C**, **Fig S5** and **Fig S6**. For analyses of siRNA relative abundances or siRNA coverage across siRNA locus category across siRNA transcriptomes, a linear model was fitted using logit transformation to correct for heteroscedasticity (**Fig 1C**, **Fig 2B**, **Fig 3A**, **Fig 3B**, **Fig 3D**, **Fig 4B, Fig S1A**, **Fig S2B**, **Fig S2D**, and **Fig S2F**). For analyses of siRNA counts or locus counts, a linear model was fitted using log(RPM + 1) transformation to correct for heteroscedasticity (**Fig 2E**, **Fig S2G**, **Fig S3A**). For analyses of distances to nearest genes, a generalized linear model was fitted using log link function to correct for heteroscedasticity (**Fig 5A**). For analyses of fraction of locus length covered by genes, a generalized linear model of quasibinomial family with logit link function was fitted to accommodate the mean-error relationship of fractional data (**Fig 2D**, **FigS3B**, **Fig 5B**). For analyses of DNA methylation levels across different locus categories, a generalized linear model of quasibinomial family with logit link function was fitted to accommodate the mean-error relationship of proportion data (**Fig 5C**, **Fig S5**, **Fig S6**). For analysis of correlations between PC1 (**Fig 4D**), distance to nearest genes, TE overlaps and DNA methylation, Spearman’s rank order correlation was used (**Fig S3D-E**, **Fig S4**). P values < 2.2e-16, which is the smallest positive floating point number R can display (R Core Team 2020), were treated as 0 by R, and reported as such in this study.

## Supplemental information

**Additional file 1: Table S1.** General mapping statistics

**Additional file 2: Table S2.** Pollination success rates

**Additional file 3: Supplemental figures.** Fig S1-S6

**Additional file 4: Supplemental Dataset1.** miRNA read counts

## Acknowledgements

We thank Zachary Liechty and Christian Santos for assistance in R programming; and Alina Yalda, Jake Anichowski, and Michelle Binyu Cui for greenhouse maintenance and technical assistance. The UC Davis Genome Center provided Illumina sequencing, library quality control and size selection services. CL also acknowledges partial support by Elsie Taylor Stocking Memorial Fellowship from the Department of Plant Biology at University of California, Davis. This study was supported in part by resources and technical expertise from the Georgia Advanced Computing Resource Center, a partnership between the University of Georgia’s Office of the Vice President for Research and Office of the Vice President for Information Technology.

## Declarations

### Availability of data and materials

All small RNA data have been deposited in the Sequence Read Archive, BioProject PRJNA533115. All R codes regarding data visualization and statistical analyses were deposited in https://github.com/cxli233/zygote_smRNA/

### Author contributions

CL, JIG, SDR and VS designed the study. HX and HF collected zygotes. SDR supervised zygote collections. CL produced small RNA sequencing libraries. CL and JIG analyzed data. VS supervised data collection and analyses. CL wrote the manuscript with input from all authors.

### Ethics approval and consent to participate

Not applicable

### Consent for publication

Not application

