## Supplemental Figures for "Resetting of the 24-nt siRNA landscape in rice zygotes"

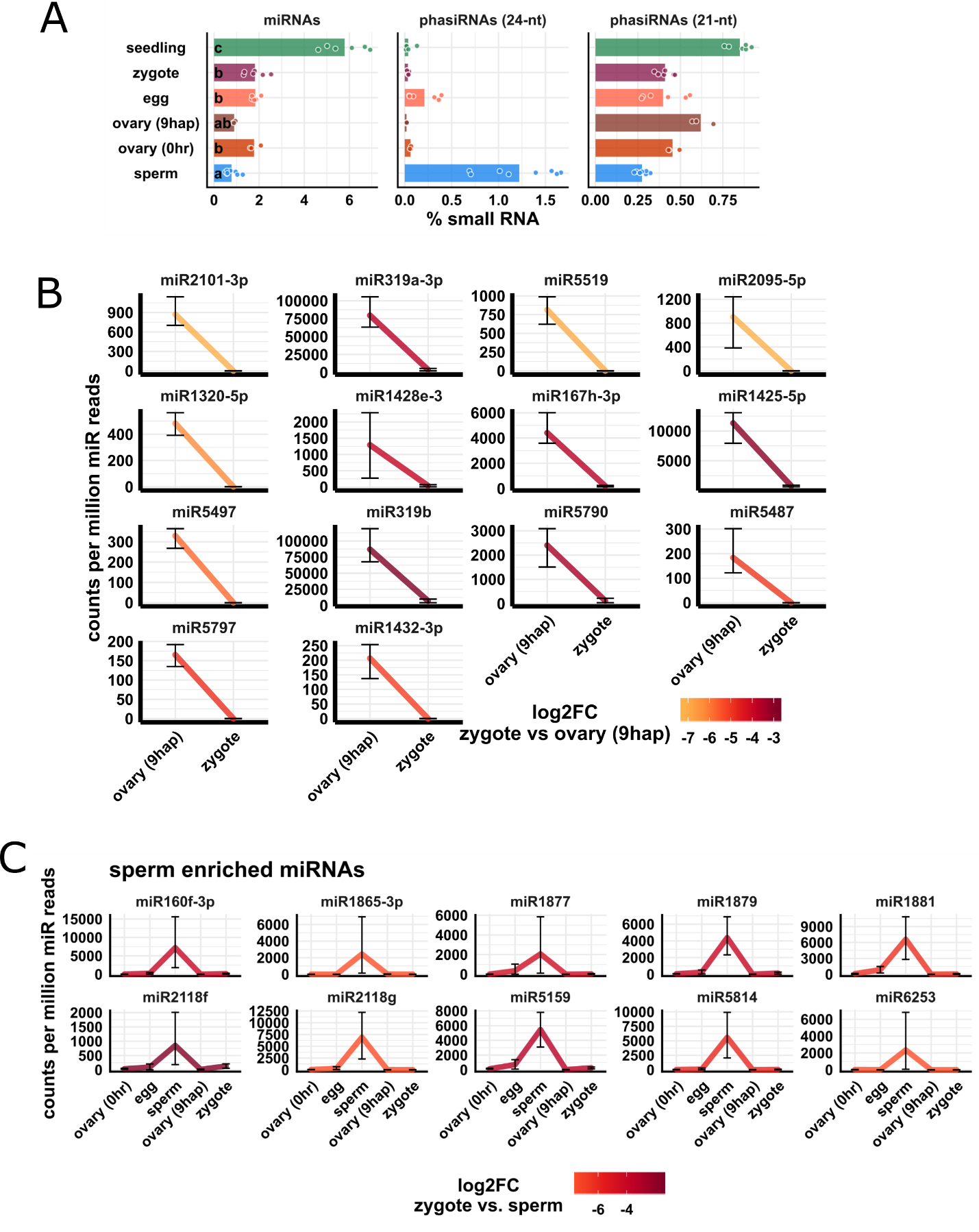

**Fig S1, supporting Fig 1**

(A) Relative abundances of miRNAs, 21- and 22-nt phasiRNAs in zygote small RNA transcriptomes. x-axis values are relative to total 20 – 25-nt small RNA reads. Each data point is a small RNA transcriptome. Letter grouping (α *=* 0.05) is based on linear models with logit transformation followed by Tukey tests.

(B) Significantly downregulated miRNAs in zygote relative to post-fertilization ovary. Differential abundance is determined by a 2-fold decrease and FDR < 0.05 cutoffs. y-axis values are relative to per million miRNA reads in each sample. Error bars are 95% confidence intervals.

(C) Additional sperm-enriched miRNAs are downregulated in the zygote (see also **Fig 1E**). Sperm-enriched is determined by >1000 reads per million miRNA reads in sperm, but < 500 reads per million miRNA reads in egg. y-axis values are relative to per million miRNA reads. Color code reflects log2FC values for zygote - sperm. Error bars are 95% confidence intervals.

Zygote and 9 hap ovary data are from this study, all other data from Li et al., (2020).

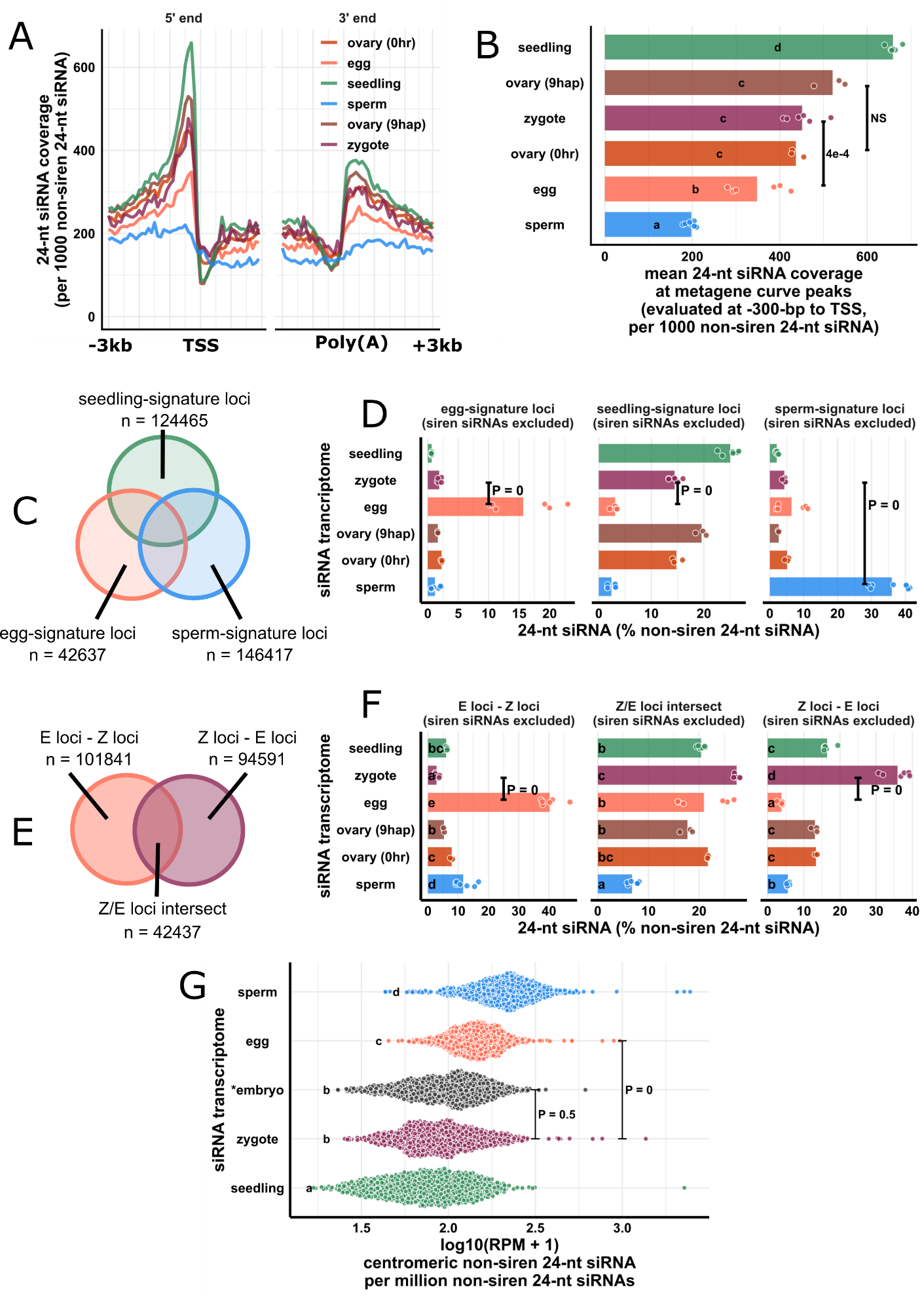

**Fig S2, supporting Fig 3 & Fig 4**

(A) Metagene coverage plot for 24-nt non-egg-siren siRNAs. Coverage is measured over 100-bp intervals and normalized per 1000 24-nt non-egg-siren siRNAs. Vertical grid lines are 500-bp intervals. TSS transcription start site, poly(A) polyadenylation site.

(B) Quantification of (A) at the interval from 300 to 200-bp upstream of TSS, corresponding to the peaks of metagene curves. Each data point is an siRNA transcriptome and bar heights are averages. x-axis values are normalized per 1000 24-nt non-egg-siren siRNAs.

(C) Venn diagram illustrating egg-signature loci (egg – seedling – sperm siRNA loci), seedling-signature loci (seedling – egg – sperm siRNA loci), and sperm-signature loci (sperm – egg – seedling siRNA loci), as in **Fig 3C**.

(D) Bar plot showing relative abundances of 24-nt siRNA across siRNA loci categories defined in **D**. The zygote siRNA transcriptome was not used to define these locus categories. Each data point is an siRNA transcriptome. Bar heights are averages. x-axis values are normalized to total 24-nt non-egg-siren siRNAs.

(E) Venn diagram illustrating E loci – Z loci, Z loci – E loci, and Z/E loci intersect, as in **Fig 4A**.

(F) Quantification of 24-nt siRNA relative abundances across siRNA loci categories defined in **F**. Each data point is a siRNA transcriptome. Bar heights are averages. x-axis-values are relative to total 24-nt non-egg-siren siRNAs.

(G) Quantification of non-siren 24-nt siRNA at centromeric regions. Each data point is a 50-kb window at centromeric regions across 12 rice chromosomes. x-axis values normalized to per million total non-siren 24-nt siRNAs and log10 transformed. Biological replicates were averaged prior to the analysis.

Letter grouping (α *=* 0.05), and P values are based on Tukey tests. Sizes of overlap in Venn diagrams are not to scale. Zygote and 9 hap ovary data are from this study; Embryo data from Rodrigues et al. (2013); all other data from Li et al. (2020).

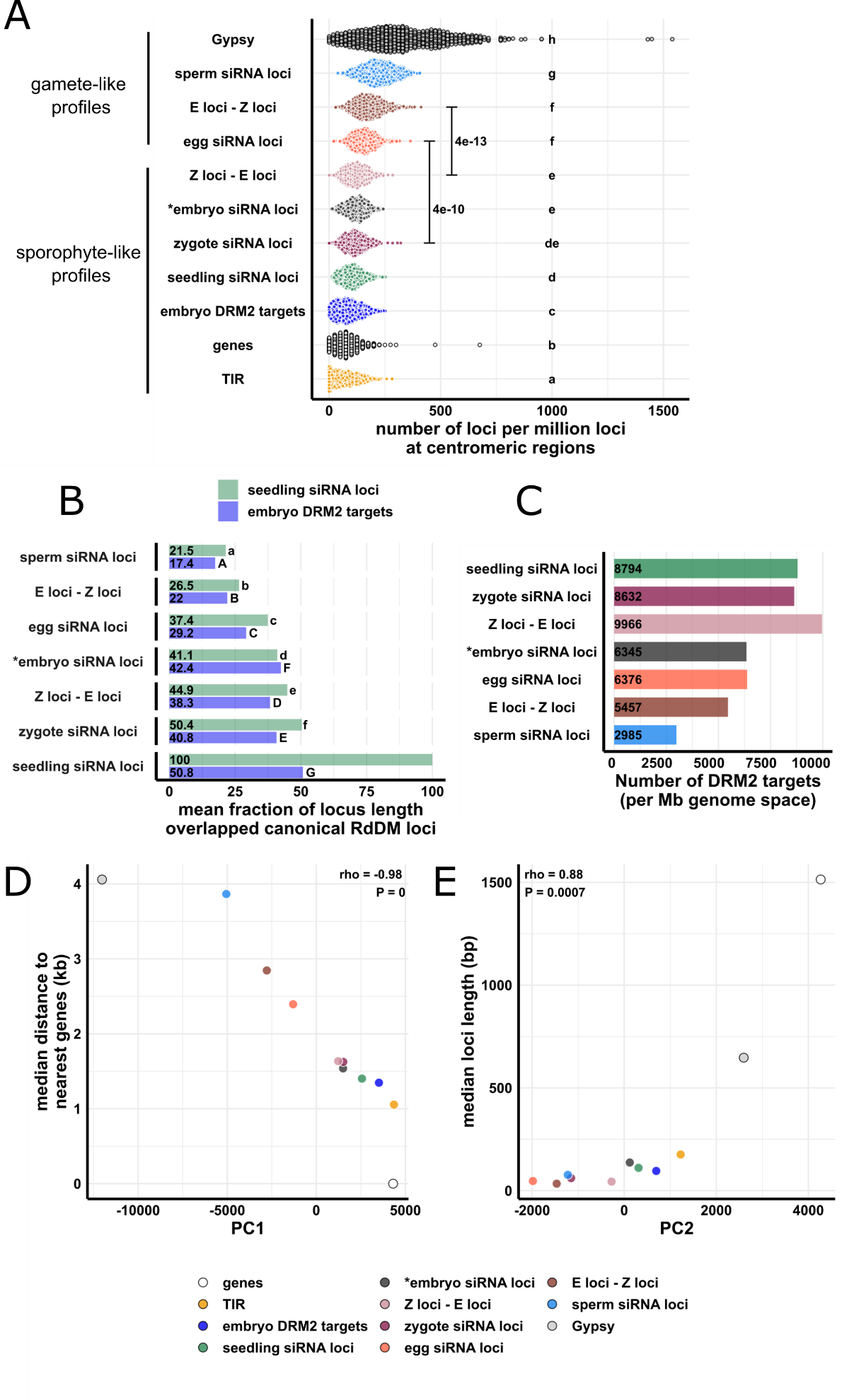

**Fig S3: supporting Fig 4 and Fig 5.**

(A) Quantification of **Fig 4C**. Each data point is a 50-kb genomic window inside centromeric regions for all 12 rice chromosomes. x-axis values are number of loci per million total siRNA loci for each loci category.

(B) Bar plots showing mean locus length overlapped by seedling siRNA or embryo DRM2 targets across siRNA loci categories. Statistical comparisons are made across siRNA loci categories within a locus category.

(C) Bar plots showing number of embryo DRM2 targets overlapped by different siRNA loci categories. x-axis values normalized to per million basepairs occupied by siRNA loci categories.

(D) Scatter plot showing correlation of PC1 (**Fig 4D**) and median distance to nearest genes (**Fig 5A**). Median distance to nearest genes for genes is set to 0.

(E) Scatter plot showing correlation of PC2 (**Fig 4D**) and median length of locus.

Letter grouping (α = 0.05) and P values are based on Tukey tests. Rho, Spearman’s rank order correlation coefficient. *Embryo siRNA data from Rodrigues et al (2013), which was based on a single replicate. Except zygote, all other data from Li et al. (2020).

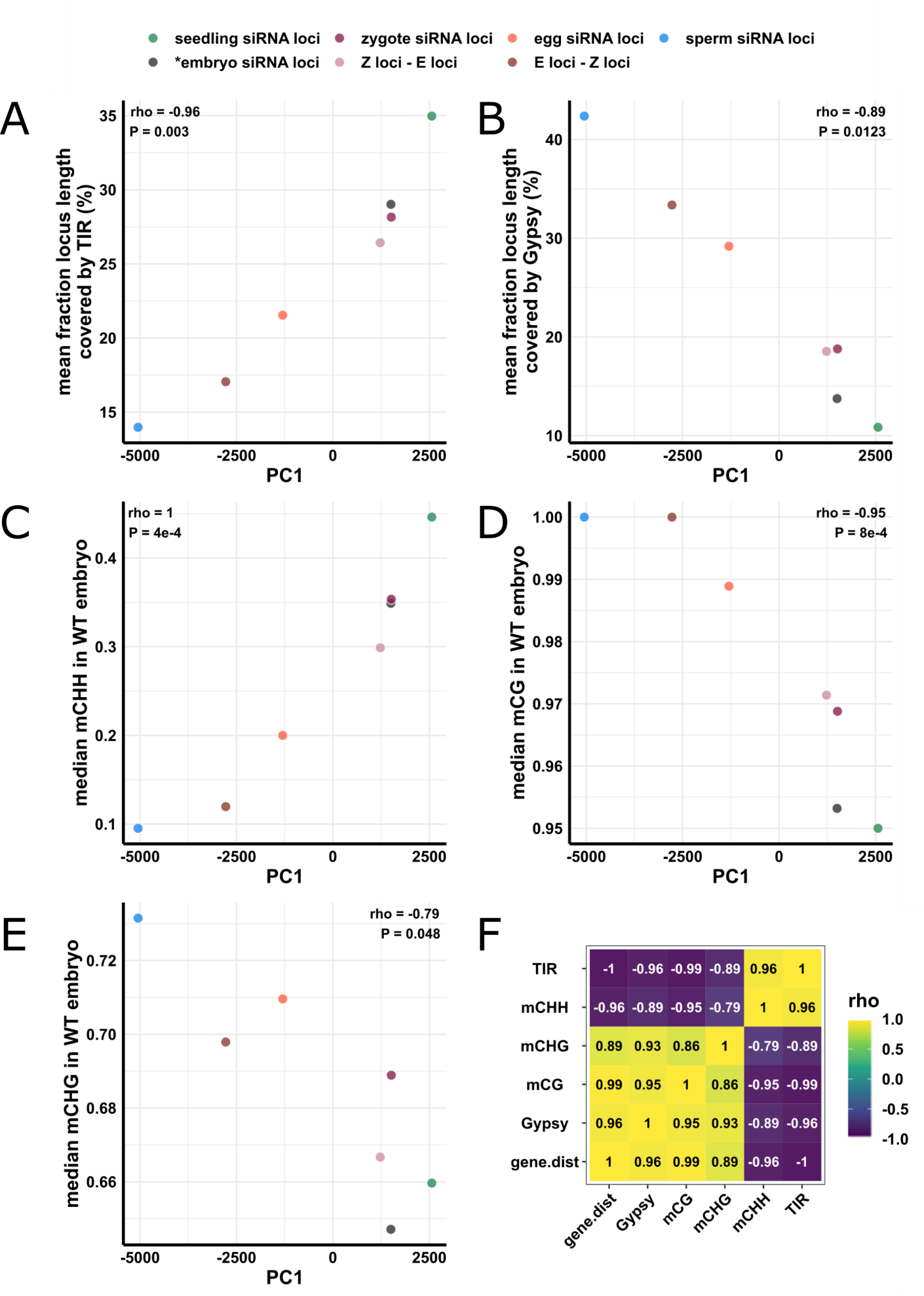

**Fig S4, supporting Fig 4 and Fig 5**

(A) Scatter plot showing correlation of PC1 (**Fig 4C**) and mean fraction of locus length covered by TIR transposon (**Fig 5B**).

(B) Scatter plot showing correlation of PC1 (**Fig 4C**) and mean fraction of locus length covered by Gypsy retrotransposon (**Fig 5B**).

(C), (D) and (E) Scatter plot showing correlation of PC1 (**Fig 4C**) and median methylation at mCHH, mCG and mCHG contexts, respectively (**Fig 5C,** see also **Fig S5**).

(F) Correlation heat map showing pairwise rank order correlation between distance to nearest genes (gene.dist, **Fig 5A**), *Gypsy* retrotransponson overlaps (**Fig 5B**), DNA methylation (**Fig 5C**, see also **Fig S5**), and TIR transposon overlaps (**Fig 5B**).

Rho, Spearman’s rank order correlation coefficient. *Embryo siRNA data from Rodrigues et al (2013), which was based on a single replicate. Except zygote, all other data from Li et al. (2020).

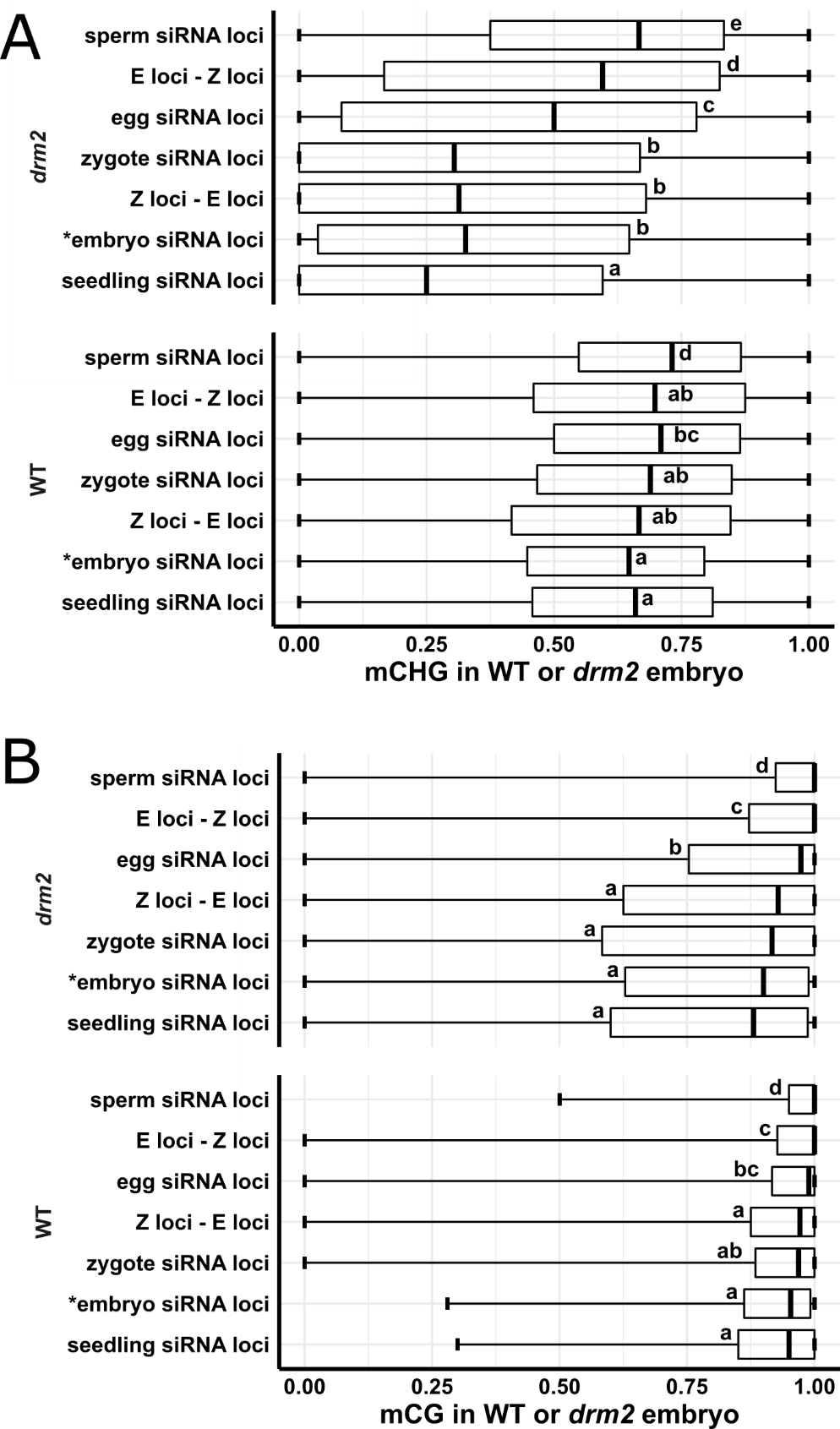

**Fig S5, supporting Fig 5**

Boxplots showing mCG (A) and mCHG (B) methylation level in wildtype or *drm2* embryo. Middle lines are median. Boxes span interquartile range. Whiskers span 2.5^th^ and 97.5^th^ percentiles. Letter groupings are based on Tukey tests. *Embryo siRNA data from Rodrigues et al (2013), which was based on a single replicate. Except zygote, all other data from Li et al. (2020).

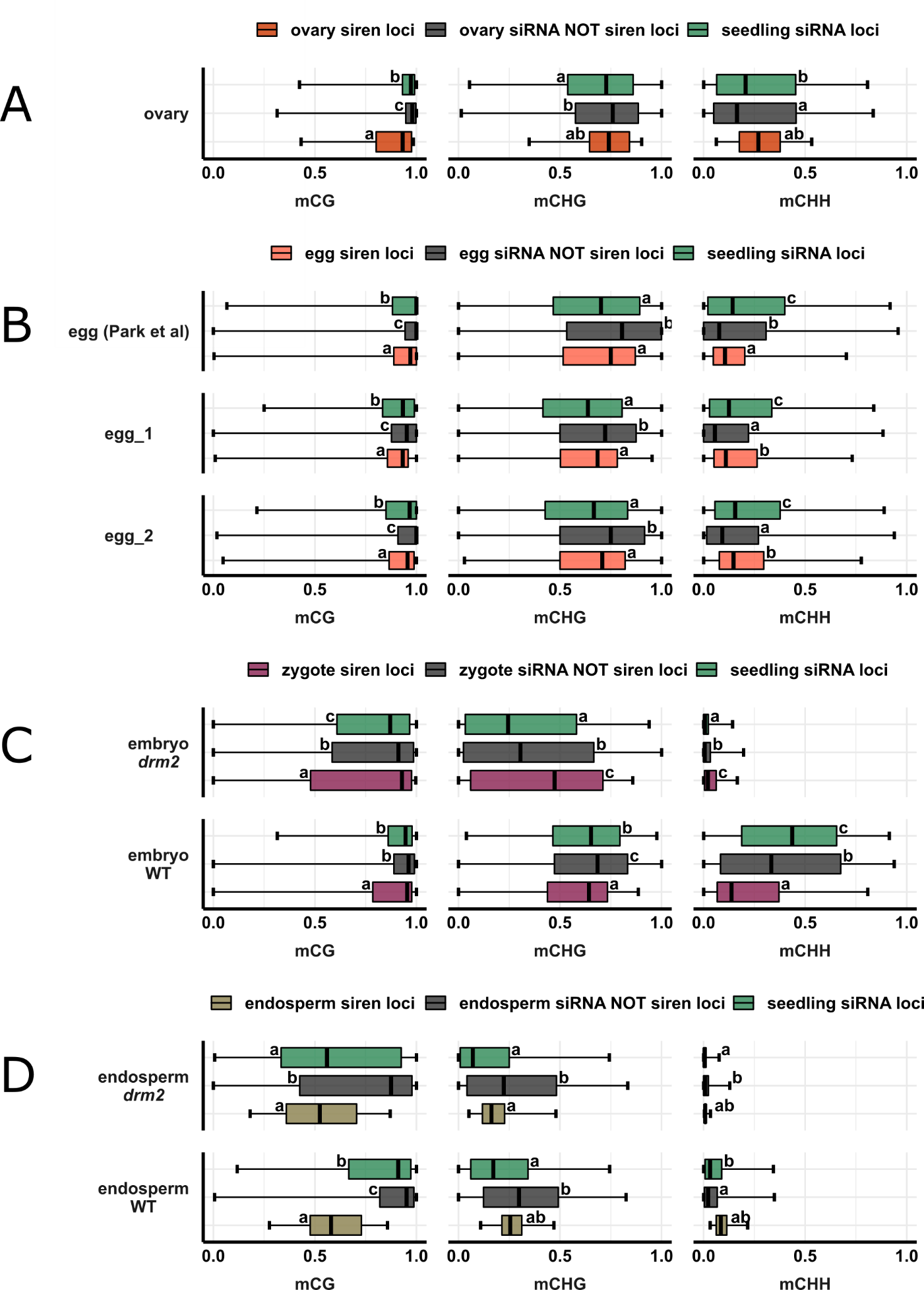

**Fig S6, supporting Fig 2 and Fig 5**

Boxplots showing DNA methylation level in ovary (A), egg cell (B), wildtype or *drm2* embryo (C) and wildtype or *drm2* endosperm (D). Middle lines are median. Boxes span interquartile range. Whiskers span 2.5^th^ and 97.5^th^ percentiles. siRNA NOT siren loci refer to siRNA loci that are not siren loci. Letter groupings are based on Tukey tests. Egg (Park et al) data from Park et al., (2016). Except zygote, all other data from Li et al. (2020).
